# Rate Limiting Enzymes in Nucleotide Metabolism Synchronize Nucleotide Biosynthesis and Chromatin Formation

**DOI:** 10.1101/2025.05.21.655426

**Authors:** Shashank Srivastava, Daniela Samaniego-Castruita, Sakshi Khurana, Vipul Shukla, Issam Ben-Sahra, Daniel R. Foltz

## Abstract

Chromatin formation requires both an adequate nucleotide supply and sufficient availability of histones. Chromatin must be assembled following DNA-based fundamental cellular processes such as replication and transcription to preserve genome integrity. Chromatin assembly is regulated by the orderly engagement of histones with a series of histone chaperones that guide newly synthesized histones from ribosomes to DNA. Although the synthesis of nucleotides and the histone proteins are the two major biosynthetic processes that complete the formation of chromatin, how these processes are coordinated remains unknown. Phosphoribosyl pyrophosphate synthetases (PRPSs) catalyze the first and rate-limiting step in the nucleotide biosynthesis pathway. PRPS enzymes form a complex with PRPS-associated proteins (PRPSAPs). In the present study, we discover that PRPS-PRPSAP enzyme complex are part of histone chaperone network. We show that PRPS enzymes are essential not only for nucleotide biogenesis, but together with PRPSAP also play a key role in the early steps in the process of histone maturation by regulating the interaction of histone chaperones with histone H3 and H4. Importantly, this regulation is separate from PRPS nucleotide biosynthetic activity. Depletion of PRPS proteins leads to limited histone availability and impaired chromatin assembly. Our discovery bridges cellular metabolism and chromatin regulation and provides the evidence of how nucleotide biogenesis and histone deposition are coordinated.

## Introduction

Chromatin formation is central to organization and spatiotemporal regulation of genome. The structural unit of chromatin is the nucleosome which consists of 147 base pairs DNA wrapped around an octamer of histone proteins: two dimers of histone H2A and H2B and a tetramer of histone H3 and H4^1–3^. Nucleosome subunits are separated by linker histone H1, allowing higher order folding and compaction of chromatin^4^. Vital cellular processes such as replication, transcription, DNA repair, which all use DNA as a template, necessitate the eviction of histones to allow for access of process-specific factors^5–8^. Immediate restoration of chromatin via redeposition of histones is pivotal to maintain genome stability and preserving epigenetic information^9,10^. Defects in chromatin assembly leave DNA vulnerable to damage and elevate genomic instability^11,12^.

Chromatin assembly is governed by a network of histone chaperones that safeguard, fold and deposit nascent histone proteins onto DNA to form nucleosomes^13,14^. Despite some ambiguity as to the order of chaperones that engage with nascent histones, according to most accepted model, the newly synthesized histone H3 and H4 as they exit ribosome, first associate with heat shock proteins HSP70/90 and histone chaperone NASP that facilitate the proper folding of histones and allow dimerization of H3-H4^15^. The growing polypeptide chain of histone H3 is also monomethylated by SetDB1 at Lys-9^16^. Histone H3 and H4 then interact with HAT1-RBBP7 complex which acetylate histone H4 at Lys-5 and Lys-12^17,18^. The posttranslational modifications H3K9me1, H4K5ac and H4K12ac are considered as bona-fide marks for new histone H3 and H4 respectively^19^. Histone H3 and H4 are then handed to ASF1 and importin-4 before engaging with replication or transcription specific chaperones, facilitating H3-H4 dimer translocation to the nucleus to assemble chromatin^15,19–21^.

Nucleotides are obligatory precursors for nucleic acid synthesis. The rate-limiting step in nucleotide biosynthesis pathway is regulated by phosphoribosyl pyrophosphate synthetases (PRPS) that catalyze the synthesis of phosphoribosyl pyrophosphate (PRPP) from ribose-5-phosphate (R5P), provided by the pentose phosphate pathway (PPP)^22,23^. PRPP is an activated sugar that through a series of tightly regulated biochemical reactions produces purine and pyrimidine nucleotides^24^. Eukaryotes have three PRPS isoforms, PRPS1, -2 and -3, however only PRPS1 and -2 are ubiquitously expressed^25^. With approximately 95% sequence similarity, PRPS1 and -2 are functionally redundant^26^. PRPSAPs have evolved from PRPS in such a way that their catalytic domain is disrupted by insertion of a new domain, non-homologous region (NHR)^27,28^. PRPS and PRPSAP together can be considered as the PRPP-synthetase ^22,23^. As a component of PRPP-synthetase enzyme complex, PRPSAPs have been shown to negatively regulate PRPS enzymes^28,29^, nonetheless, their exact function in nucleotide biosynthesis and beyond remain largely unknown.

Sufficient supply of nucleotides is critical for DNA synthesis^30^. An acute drop in nucleotide levels can induce replication stress potentially causing replication fork collapse and cell death^12,31^. Likewise, insufficient histone supply, perturbations in histone chaperone pathways and histone oversupply lead to cell cycle arrest and increased genome instability^32,33^. Nucleosomes are immediately assembled following DNA replication using both old (parental) and new histones, and in this manner replication coupled nucleosome assembly provides an elegant and efficient way to maintain genome stability and ensure epigenetic inheritance^8,34^.

Cells depend on nucleotide availability and nucleosome assembly concurrently for duplication of the genome, suggesting there may be a coordinated mechanism synchronizing nucleotide biosynthesis with histone deposition. This synchrony remains unexplored, yet such a synchrony would ensure tightly orchestrated nucleotide production and replication-coupled nucleosome assembly, leading to precision in both genome duplication and chromatin formation.

Here, we discover a novel role for PRPS-PRPSAP in chromatin assembly, distinct from the well-established function of PRPS in nucleotide biosynthesis. This finding reveals that PRPS enzymes have evolved to perform dual functions, driving nucleotide production while also facilitating histone deposition during chromatin assembly. Our study establishes PRPS as a key regulator that links nucleotide metabolism to chromatin assembly, shedding light on the molecular mechanisms that coordinate these essential cellular processes.

## Results

### PRPS loss leads to an accumulation in newly synthesized histone marks

To gain insight into how nucleotide metabolism and histone deposition are coordinated, we perturbed the rate-limiting step of nucleotide biosynthesis by knocking down the PRPS1 or PRPS2 enzymes. These enzymes control this crucial step in the pathway and are functionally redundant (Figure 1A). We examined whether this disruption affects histone H3 and H4 levels, along with H3K9me1, H4K5 and K12ac. These posttranslational modifications are associated exclusively with newly synthesized histone H3 and H4 and acquired prior to histone deposition into the chromatin (Figure 1A). We observed an increase in steady-state levels of both histone H3 and H4 upon PRPS1 or PRPS2 knockdown (Figures 1B, S1A). The increase in histones H3 and H4 was accompanied by an elevated H3K9me1, H4K5 and K12ac marks (Figures 1B, S1A). These data suggest that loss of PRPS1 or PRPS2 results in an accumulation of newly synthesized histones H3 and H4, indicating a potential role for PRPS enzymes in histone deposition.

**Figure 1.**
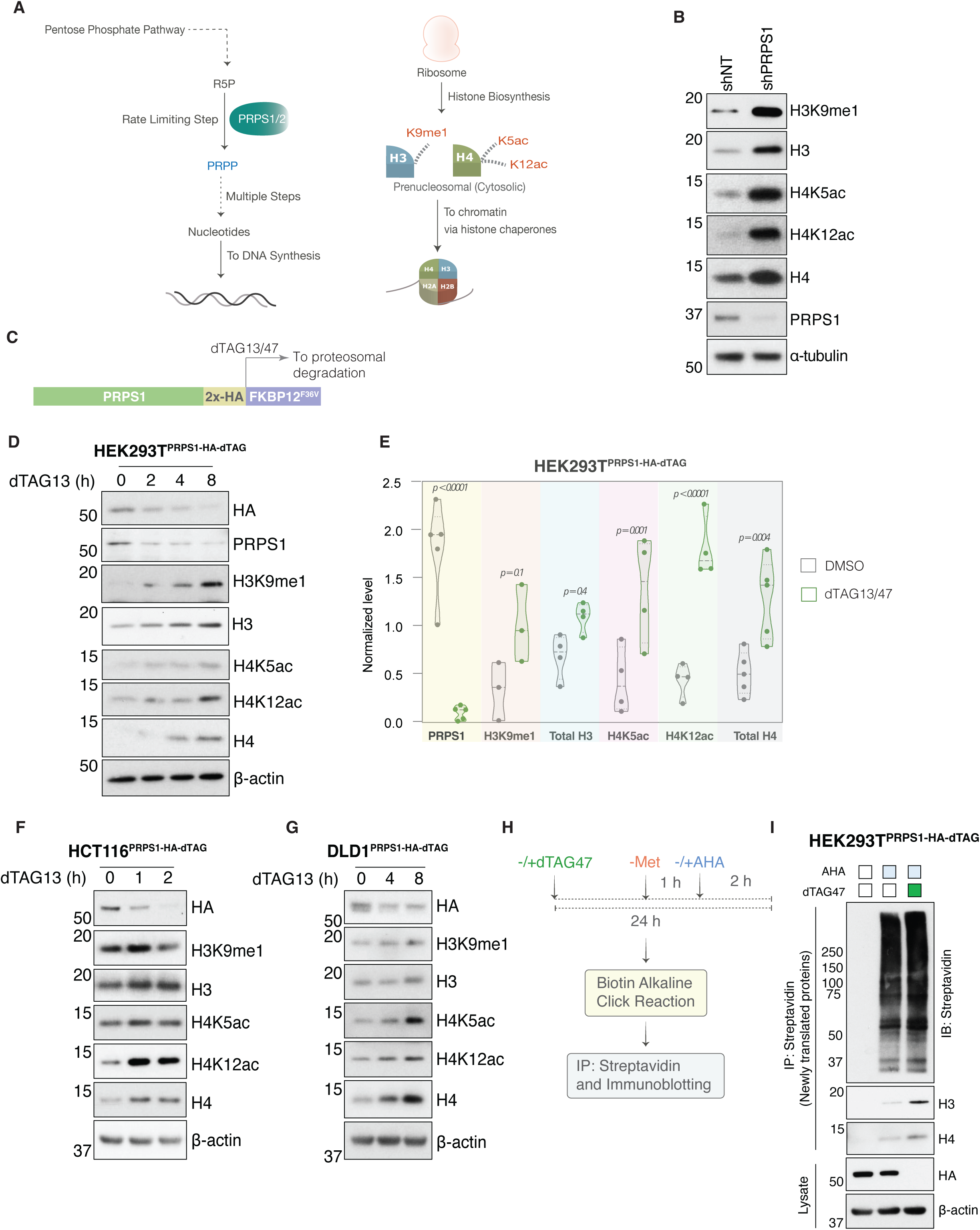
PRPS loss leads an accumulation of newly synthesized histones. **(A)** Model showing the PRPS1/2 enzymes as key regulators of nucleotide biosynthesis (left) and histone deposition pathway from histone biosynthesis with posttranslational marks on new histone H3 and H4 (right). **(B)** Total histone and posttranslational modification marks associated with new histones in Ctrl or PRPS1 shRNA treated HEK293T cells. **(C)** PRPS1-HA-FKBP12^F36V^ (dTAG) schematic and induction of its proteasomal degradation in response to dTAG13/47. **(D)** Total histone and posttranslational modification marks associated with new histones in response to time dependent depletion of PPRS1 in HEK293T^PRPS1-HA-dTAG^ cells. **(E)** Violin plot below show the levels of PRPS1-HA-dTAG, total histone H3, H4 and posttranslational modification marks associated with new H3 and H4, normalized to β-actin from multiple independent experiments in HEK293T-PRPS1-HA-dTAG cells treated with dTAG13 or 47 for 24 hours. *p* values were calculated using two-way ANOVA. **(F-G)** Total histone and posttranslational modification marks associated with new histones in response to time dependent depletion of PPRS1 in HCT116^PRPS1-HA-dTAG^ *(F)* or DLD1^PRPS1-HA-dTAG^ *(G)* cells. **(H)** Experimental workflow for labeling newly synthesized proteins with AHA and subsequent isolation through streptavidin immunoprecipitation. **(I)** Histone H3 and H4 in the pool of newly synthesized protein isolated from HEK293T^PRPS1-HA-dATG^ cells with or without PRPS1 depletion. PRPS1 depletion is shown in whole cell lysate.

shRNA mediated knockdowns are not acute, limiting their ability to specifically assess the biological effects resulting from the loss of a protein. To evaluate the role of PRPS enzymes in histone deposition we engineered cell lines with FKBP12^F36V^ degron-tag at endogenous PRPS1 and -2 loci in HEK293T, HCT116 and DLD1 cells (Figures 1C, S1B, C). The heterobifunctional molecule, dTAG (dTAG13 or dTAG47) engages Cereblon E3 ligase complex to FKBP12^F36V^ to induce the rapid proteasomal degradation of FKBP12^F36V^ fusion chimera^35^. PRPS1-HA-FKBP12F36V (PRPS1-HA-dTAG hereafter) began to decay within 2 hours of dTAG13 treatment in HEK293T cells and by 8 hours, considerable degradation of PRPS1 was observed in HEK293T cells (Figure 1D). The rapid depletion of PRPS1 was associated with an increase in total histone H3 and H4 levels, as well as post-translational modifications H3K9me1, H4K5ac, and H4K12ac (Figure 1D). The elevated total histone H3, H4, and modifications linked to newly synthesized histones, were consistently observed across multiple independent experiments following 24 hours of dTAG-mediated PRPS1 depletion (Figure 1E). In HCT116 and DLD1 cells also, dTAG13 mediated PRPS1 depletion led to a time dependent increase in total histone H3, H4, and modifications associated with newly synthesized H3 and H4 (Figures 1F, G). A similar pattern for total and modified histone H3 and H4 were seen when PRPS2-V5-FKBP12^F36V^ (PRPS2-V5-dTAG from hereafter) was depleted in response to dTAG13 in HEK293T and HCT116 cells (Figure S1D).

To further confirm that PRPS1 depletion results in an accumulation of newly synthesized histones, HEK293T^PRPS1-HA-dTAG^ cells, with or without dTAG47 treatment, were first depleted of methionine and subsequently labeled with methionine analog L-azidohomoalanine (AHA)^36^. This labeling allowed tagging of newly translated proteins that were subsequently biotinylated via click chemistry and purified using streptavidin precipitation (Figure 1H). Strikingly, histones H3 and H4 were enriched in the newly translated protein pool in PRPS1-depleted cells, providing strong evidence that PRPS1 depletion leads to the accumulation of newly synthesized histones H3 and H4 (Figure 1I).

Histones are encoded by multiples genes and, the increase in histone H3 and H4 protein levels was not predominantly reflected at the transcript levels in subset of genes we examined. There were no significant changes in the transcript levels of histone H3 and H4 genes H3FA, H4C3, and H4C5. However, a moderate increase was observed in the H3F3A transcript upon PRPS1 depletion (Figure S1E).

### Elevated histone levels resulting from PRPS loss are independent of nucleotide biosynthesis pathway or cell cycle position

As PRPS enzymes are critical for nucleotide biosynthesis, we asked whether observed effects on new histone upon PRPS1 depletion result from defective nucleotide biosynthesis or indicate a nucleotide biosynthesis independent mechanism. If the accumulation in new histone following PRPS1 depletion is due to defective nucleotide biosynthesis pathway, direct inhibition of steps downstream to PRPS1 should yield similar effects on histone levels. We inhibited phosphoribosylglycinamide formyltransferase (GART), a key enzyme downstream of PRPS1 in the purine synthesis pathway, by lometrexol (LTX) under dialyzed serum conditions that restricts exogenous sources of nucleobases and nucleosides, and tested its effect on histone levels (Figure 2A). In HEK293T^PRPS1-HA-dTAG^ cells, 8-hour LTX treatment significantly reduced phospho-S6 kinase levels, used as a positive control for effective nucleotide biosynthesis inhibition^37^ (Figure 2A). Despite the inhibition of nucleotide biosynthesis, total histone H4, H4K5ac, and H4K12ac levels remained unchanged. Histone H3 and H3K9me1 showed a slight decrease, but importantly, histone levels did not increase, unlike the response to acute PRPS1 depletion (Figure 2A). In addition, PRPS1 levels did not change in response to LTX treatment (Figure 2A). Overall, these results demonstrate that the effects of PRPS1 depletion on histones involves functions beyond its role in nucleotide biosynthesis.

**Figure 2.**
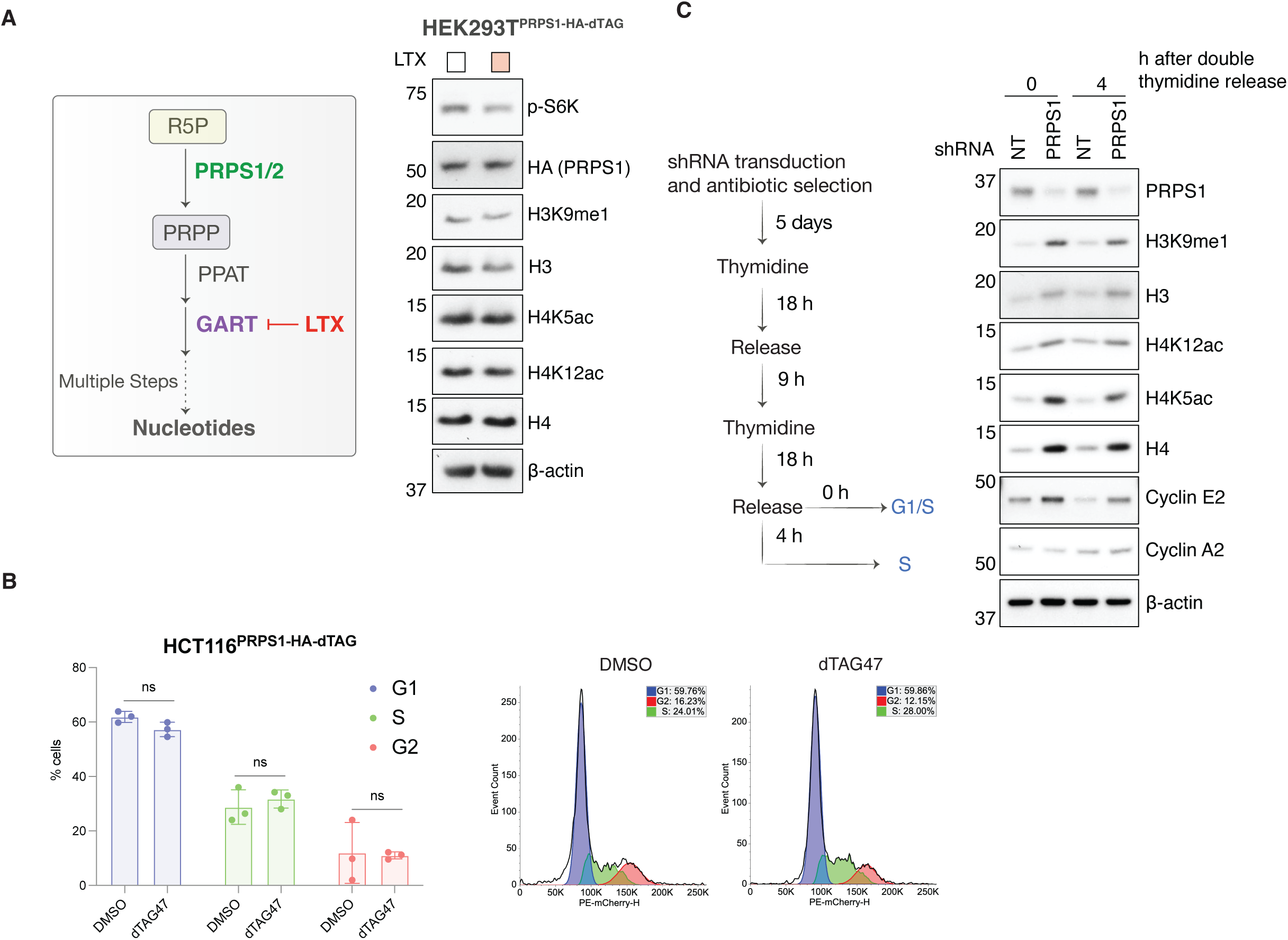
Elevated histone levels resulting from PRPS loss are independent of nucleotide availability or cell cycle position. **(A)** Schematic showing GART as a key enzyme, downstream to PRPS, in nucleotide biosynthesis and its inhibition from LTX. *(left)*. PRPS1 (HA), total histone and posttranslational modification marks associated with new histones in HEK293T^PRPS1-HA-dTAG^ cells treated with or without 1 μM LTX for 8 h. pS6K was used as a positive control for nucleotide synthesis inhibition *(right)*. **(B)** Cell cycle profile of HCT116^PRPS1-HA-dTAG^ cells treated with or without dTAG13 for 24 h, showing the distribution across G1, S and G2 phases. Mean and standard deviation is shown from three independent experiments in bar plot, histogram is representative of one of the experiments. **(C)** Immunoblot showing the histones in HEK293T cells transduced with Ctrl. or PRPS1 shRNA and synchronized with double thymidine block, as depicted in the schematic to the left of the blot.

PRPS1 deficiency leads to cell cycle arrest in G1 phase due to insufficient nucleotide supply^38^ and cells with halted DNA replication would be defective in histone deposition^38^. However, the short-term effect of acute PRPS depletion on cell cycle progression is unknown. To determine whether the observed changes in histones are a direct consequence of PRPS1 depletion or a secondary effect of cell cycle arrest, we analyzed the cell cycle profiles of HCT116 cells with or without PRPS1 depletion for 24 hours. Cell cycle profiling using flow cytometry revealed that dTAG-mediated PRPS1 depletion for 24 hours does not significantly alter the G1, S, or G2/M phases in HCT116 cells (Figure 2B). This suggests that the acute elimination of a single PRPS enzyme for 24 hours is insufficient to arrest cells, and that nucleotide levels are sufficient to support DNA replication under these conditions. Moreover, the observed accumulation of histones during this period suggests that PRPS1 directly influences histones, independent of any cell cycle changes.

Histone biosynthesis begins as cells transition to S phase from G1 phase^39^, and histone deposition is accompanied with DNA replication during S-phase^34^. To test if PPRS1 loss has an impact on newly synthesized histone H3 and H4 in S-phase, control or PRPS1-shRNA cells were synchronized with double thymidine block to arrest the cells at G1/S phase boundary and released from synchrony for 4 hours to acquire mid S-phase population (Figure 2C). Consistent with our observation in asynchronous cells, PRPS1 loss resulted an accumulation of newly synthesized histone H3K9me1, H4K5 and K12 acetylation marks along with total an increase in total histone H3 and H4 in the beginning and mid S-phase (Figure 2C). This result highlights a connection between PRPS1 and histones, particularly in S-phase when histones are actively involved in nucleosome assembly.

### New histone deposition into nucleosomes depends on upstream PRPS function

After synthesis, histones are rapidly translocated to the nucleus through histone chaperone pathway to assemble chromatin^13,14^. Given the elevated levels of new histone H3 and H4 marks, we posited a role for PRPS in histone deposition to form chromatin. To explore this idea, we first examined the levels of histones in soluble cytosolic and endonuclease (Benzonase) digested chromatin fraction (Figure 3A). In HEK293T and DLD1 cells, dTAG47 mediated PRPS1 depletion resulted in an increased newly synthesized histone marks H3K9me1, H4K5 and K12ac along with total histone H3 and H4 in the soluble pre-nucleosome fraction (Figure 3B). In contrast, new histone marks and total histone H3 and H4, were diminished in chromatin fraction upon PRPS1 depletion (Figure 3B). New histone marks persist for a short time in chromatin following nucleosome deposition, and therefore, their presence in chromatin is a marker of new histone deposition ^40^. Additionally, we conducted an unbiased epiproteomic approach to profile the histone posttranslational modifications present in chromatin of PRPS1 or -2 knockdown HEK293T cells (Supplementary Table S1). This approach also revealed a striking decrease in H4K5 and K12 acetyl modified histones in the chromatin upon PRPS1 or PRPS2 loss (Figure S2A). H3K9me1 decrease was observed only with PRPS1 knockdown (Figure S2A).

**Figure 3.**
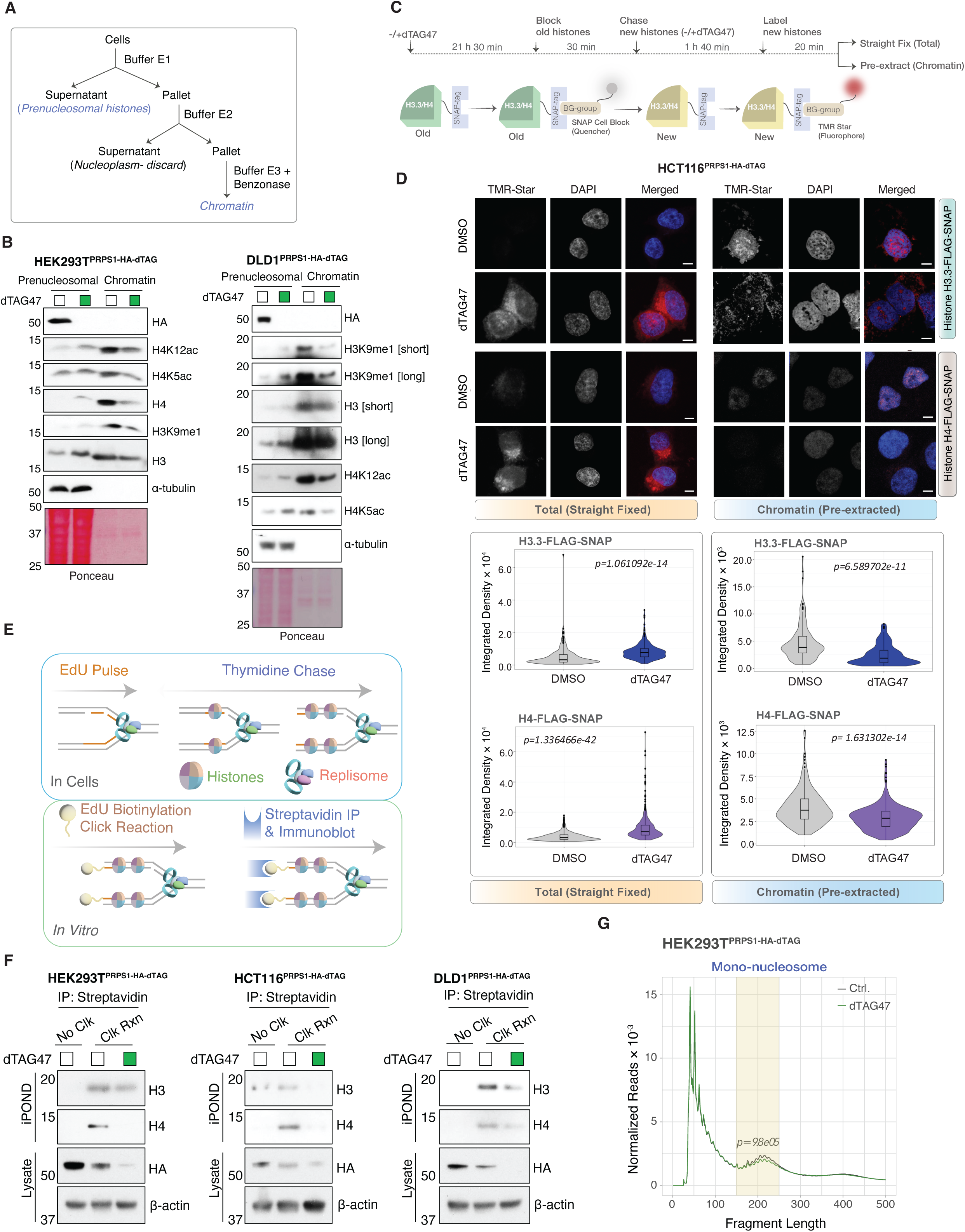
PRPS1 depletion causes new histone deposition impairment. **(A)** Schematic showing the isolation of prenucleosomal and chromatin bound histones through benzonase digestion **(B)** Immunoblot showing the total histone and posttranslational modification marks associated with new histones from the prenucleosomal or nucleosome-incorporated histones from the chromatin fraction of HEK293T^PRPS1-HA-dTAG^ and DLD1^PRPS1-HA-dTAG^ cells treated with DMSO or dTAG47 for 24 h. **(C)** Schematic and experimental workflow showing histone-SNAP tag system in HCT116^PRPS1-HA-dTAG^ cells to examine the new histone H3.3 and H4. (**D)** Distribution of total (straight fixed) or chromatin bound (pre-extracted) histone H3.3 or H4 form HCT116^PRPS1-HA-dTAG^ ^H3.3/H4-FLAG-SNAP^ cells with or without PRPS1 depletion. Pseudo-colors were used in TMR-Star and DAPI panels to enhance contrast, *scale bar: 5 μm*. Violin plot below represents the quantification from n=3 independent experiments. *p* values were calculated from Student’s t-test. See *Figure S3D* for orthogonal projections showing TMR-Star and DAPI non-overlapping signal in straight fixed condition. **(E)** Schematic showing the workflow of isolation of proteins from nascent chromatin **(F)** Immunoblot below showing the histone H3 and H4 associated with newly replicated DNA in HCT116^PRPS1-HA-dTAG^ and DLD1^PRPS1-HA-dTAG^ cells with or without PRPS1 depletion. **(G)** Fragment length distribution from nucleosome positioning analysis of ATAC-seq signal from HEK293T^PRPS1-HA-dTAG^ cells treated with DMSO or dTAG47 for 24 hours. Fragment lengths (150-250 bp) corresponding to mononucleosome is highlighted. *p* value was computed using Wilcoxon-singed rank test, *n =2*.

To specifically test the effect of PRPS1 loss on histone deposition, we employed covalent-fluorescent pulse chase labeling approach by SNAP-tagging histone H3.3 (DNA replication independent H3 variant) and H4 in HCT116^PRPS1-HA-dTAG^ background (Figures S2B, C). SNAP-tag is a mutant version of the DNA repair protein O^6^-alkylguanine-DNA alkyltransferase that specifically reacts with substrates derivatives of benzylguanine resulting in covalent binding of SNAP-tag with a fluorescent probe^41,42^. The preexisting SNAP-tagged histone H3.3 or H4 were quenched using non-fluorescent SNAP-cell block followed by pulse labeling of newly synthesized SNAP-tagged histone H3.3 or H4 with the fluorescent substrate TMR star in HCT116 cells with unperturbed or depleted PRPS1 (Figure 3C). We chased TMR labeled SNAP-histone H3.3 or H4 for 2 hours (Figure 3C), a timeframe sufficient for histone deposition undergoing S-phase or through transcription or DNA-mediated chromatin assembly^41^. Cells were either straight fixed to measure the total histone levels, or pre-extracted to retain only chromatin bound histones (Figure 3C). Consistent with our biochemical assays showing the accumulation of new histone marks, including H4K5ac and H4K12ac, newly synthesized SNAP histone H3.3 and H4 accumulate in the cytoplasm upon PRPS1 depletion (straight fixed, Figures 3D, S2D). Additionally, majority of histone H3.3 or H4 were seen on the chromatin with unperturbed PRPS1 within 2 hours post-labeling, whereas a significant reduction was observed in histone deposition upon PRPS1 depletion (pre-extracted, Figure 3D). H3.3 deposition reflects histone deposition into active transcription and heterochromatin outside of S-phase, and histone H4 deposition includes replication dependent and independent nucleosome assembly.

To substantiate the role for PRPS in histone deposition, we specifically measured histones on replicating DNA in presence or absence of PRPS1 through isolation of proteins on nascent chromatin (iPOND)^43^. DNA replication is a prime example where histone deposition occurs to assemble chromatin. Cells were pulse labeled with EdU to label actively replicating chromatin (Figure 3E). Biotinylation of EdU through click chemistry followed by precipitation with streptavidin allowed the identification of proteins associated with replicating DNA (Figure 3E). iPOND revealed a robust loss of histone H3 and H4 on replicating DNA upon PRPS1 depletion in three cell lines, HEK293T, HCT116 and DLD1 (Figure 3F). In addition, examining the fragment sizes generated by ATAC-seq (Assay for Transposase-Accessible Chromatin) analysis of PRPS1-depleted versus control cells revealed a significant decrease in the read counts of fragment lengths corresponding to mono-nucleosomes (150-250 bps) in PRPS1-depleted cells compared to control cells (Figure 3G), consistent with reduced nucleosome protection. Collectively, biochemical fractionation, SNAP-labeling, iPOND and ATAC experiments demonstrate that the deposition of new nucleosomes is regulated by the availability of PRPS1.

### PRPS-PRPSAP complex integrate into histone chaperone network

PRPS1 and PRPS2 enzymes are found together in a complex with PPRS-<u>a</u>ssociated <u>p</u>roteins, PRPSAP1 and PRPSAP2. While PRPS1 and PRPS2 share nearly 95% identity in their amino acid sequences, PRPSAP1 and PRPSAP2 share 75% amino acid identity with each other (Figures S3A, B). Intriguingly, PRPSAPs are also about 50% identical with PRPS enzymes (Figure S3A, B). The non-homologous region (NHR) in PRPSAPs, resulting from the disruption of the catalytic domain in ancestral PRPS, underscores their regulatory role in the PRPS enzyme complex and suggests broader functions for the PRPS-PRPSAP complex beyond nucleotide metabolism.

Given that PRPSAP is an integral part of PRPS enzyme complex, we asked if loss of PRPSAP also influences histone levels. Contrary to PPRS loss, PRPSAP1 knockdown resulted in reduced levels of newly synthesized histone H4 marks K5 and K12ac along with total histone H3 and H4 (Figure S3C). To further test the specific effect of PRPSAP1 loss, we created homozygous cell lines expressing PRPSAP1 fused with 2x-V5-BromoTag degron, replacing endogenous PRPSAP1 (Figure S3D). BromoTag is Brd4 bromodomain L387A variant that allows the selective and rapid degradation of BromoTag proteins in response to small molecule degrader AGB1^44^. In HEK293T cells, PRPSAP1-V5-BromoTag decayed within 2 hours AGB1 treatment (Figure S3E). Consistent with the results of PRPSAP1 knockdown, PRPSAP1 acute degradation also led to a rapid decrease in total histone H3 and H4 (Figure S3E). These data suggest that PRPS and PRPSAP proteins both contribute to early steps in histone deposition pathway but have distinct roles.

To gain mechanistic insight into how PRPS and PRPSAP are involved in histone deposition, we employed FLAG-affinity purification followed by mass spectrometry from HEK293T cells to uncover PRPS1, 2 and PRPSAP1 and -2 interacting partners (Supplementary Table S2 and S3). Intriguingly, this proteomic approach revealed several early cytoplasmic histone chaperones in both PRPS and PRPSAP purifications (Figures 4A, B and Figures S4A, B). The engagement of newly synthesized histones with histone chaperones is critical for histone deposition to assemble chromatin and some of the key histone chaperones enriched with PRPS and PRPSAP included HSP70 (HSPA8 and HSPA1A isoforms), HSP90 (HSP90AA1 and HSP90AB1 isoforms), which are required for proper histone folding of H3 and H4, HAT1 and RBBP7 that acetylate newly synthesized histone H4, and importin 4 (IPO4), which governs the translocation of new histone from the cytosol to the nucleus prior to chromatin assembly (Figures 4A, B). In addition, affinity purification-mass spectrometry of t- and s-isoforms of histone chaperone NASP, that promotes H3-H4 assembly^15^, revealed an enrichment of PRPS1 (Supplementary Table S4, Figures 4C, S4C). These proteomic results reveal PRPS and PRPSAP as new components of histone chaperone network and we hypothesized PRPS and PRPSAP influence histone deposition by regulating histone chaperone function.

**Figure 4.**
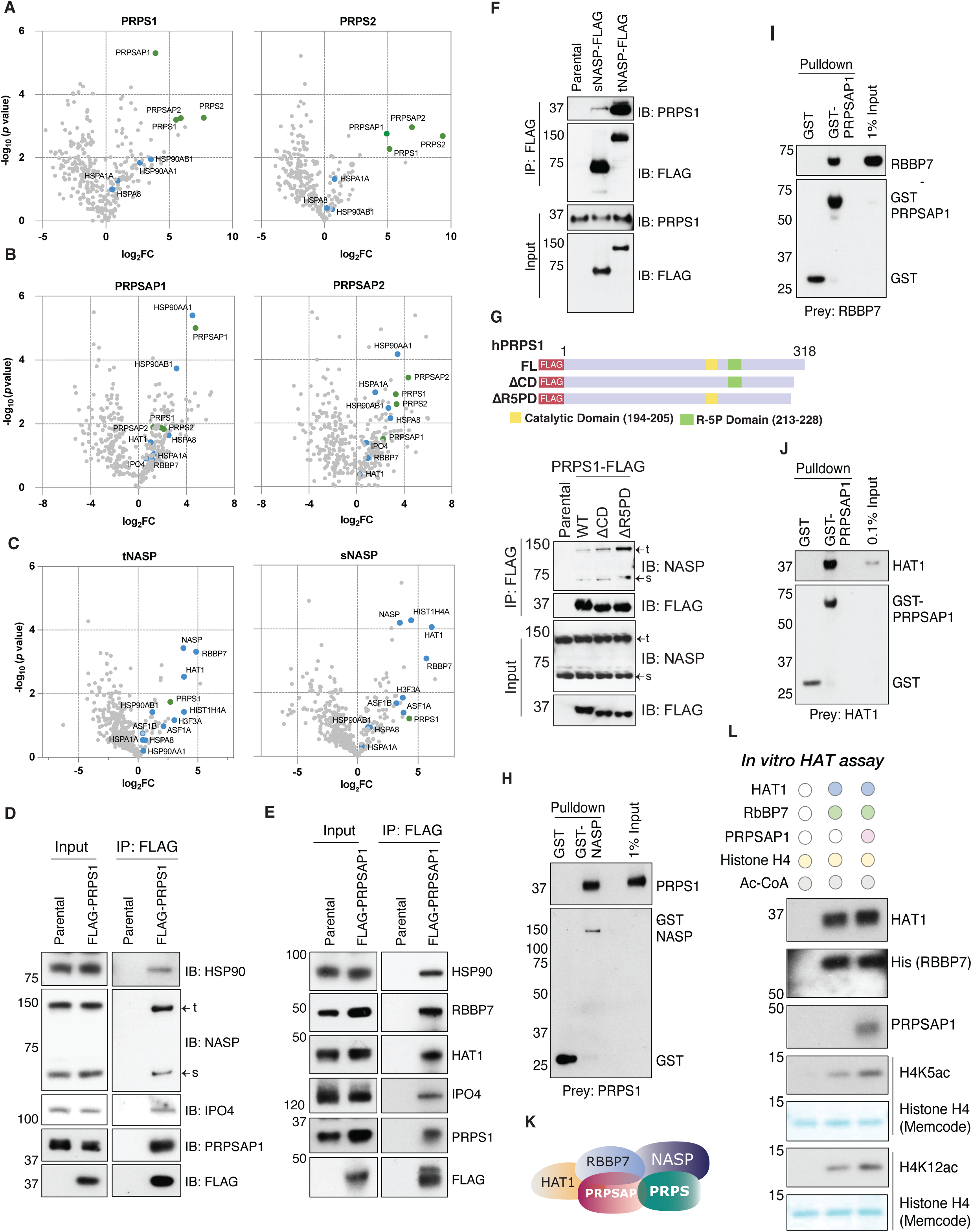
PRPS and PRPSAP integrate into histone chaperone network. **(A-C)** Volcano plots showing the proteins identified from PRPS1, PRPS2 *(A)*, PRPSAP1, PRPSAP2 *(B)*, tNASP and sNASP *(C)* FLAG-affinity purification followed by MS. PRPP-synthetase components are indicated in green whereas key histone chaperones and histones are indicated in blue color. *n=3*, *p* values were calculated using Student’s t-test with Welch correction. See materials and methods for detail. **(D-E)** Co-immunoprecipitations showing the interaction PRPS1 *(D)* or PRPSAP1 *(E)* with histone chaperones HSP90, NASP, IPO4, HAT and RBBP7 in HEK293T cells stably expressing FLAG-tagged PRPS1 *(D)* or PRPSAP1 *(E)*. **(F)** Interaction of t and s isoforms of NASP with PRPS1 as detected through co-immunoprecipitation of FLAG-tagged t- or sNASP from HEK293T cells followed by immunoblotting with PRPS1 antibody. **(G)** *Top:* Schematic showing FLAG-tagged PRPS1 wild type (WT) catalytic deficient (ΔCD) and ribose-5-phosphate binding (ΔR5P). *Below*: co-immunoprecipitation showing the interaction of FLAG-PRPS WT, ΔCD, ΔR5P with t- or sNASP in HEK293T cells. **(H-J)** GST pulldown demonstrating the in vitro binding between GST-sNASP and PRPS1 *(H)*, GST-PRPSAP1 and HAT1 *(I)* and GST-PRPSAP1 and RBBP7 *(J)*. **(K)** Depiction of PRPP-synthetase and histone chaperone complex based on the direct interaction of PRPS1 with NASP and PRPSAP1 with HAT1 and RBBP7. **(L)** *In vitro* HAT assay showing the stimulatory effect of PRPSAP1 on HAT1-RBBP7 activity towards histone H4 acetylation.

Histone chaperones HSP90, importin-4, NASP, HAT1 and RBBP7 showed a strong association with PRPS1 or PRPSAP1, through coimmunoprecipitation assays, validating our proteomic results (Figures 4D-F). Next, we tested if the catalytic domain or ribose 5-phosphate domain of PRPS1 is required for the interaction with NASP. Both catalytic deficient and ribose 5-phosphate binding mutant PRPS1 showed an enhanced interaction with NASP (Figure 4G). This implies that both catalytic and substrate binding domains of PRPS1 are not involved in the binding of histone chaperone NASP and reinforces the idea that the role for PPRS function in chromatin assembly is distinct from its role in nucleotide biosynthesis.

As affinity purification of NASP showed enrichment of PRPS1, and PRPSAP1 purification revealed enrichment of HAT1 and RBBP7, we hypothesized a direct interaction between PRPS1 and NASP, as well as between PRPSAP1 and HAT1, and RBBP7. To test this, we performed *in vitro* GST pulldown assays using recombinant GST-NASP and GST-PRPSAP1 as baits and recombinant PRPS1, HAT1 and RBBP7 as preys, which demonstrated direct binding of NASP with PRPS1, and PRPSAP1 with HAT1 and RBBP7 (Figures 4H-J). Taken together, our interaction data suggest that PRPS and PRPSAP form a complex with histone chaperones NASP, HAT1 and RBBP7 (Figure 4K).

We next investigated if PRPS or PRPSAP loss affects the levels of histone chaperones. Histone chaperones levels were examined in a time dependent manner by acute depletion of PRPS1 or PPRSAP1 by treatment with dTAG47 or AGB1, respectively. While a time dependent increase in new histone marks H3K9me1, H4K5 and K12ac were seen along with total histone H3 and H4 upon PRPS1 degradation, the HSP90, HAT1, importin-4, and NASP chaperones remained largely unchanged (Figure S4D). On the other hand, RBBP7 was increased in time dependent manner upon PRPS1 depletion, supporting increased H4K5 and K12 acetylation (Figure S4D). In contrast, PRPSAP1 resulted in a decrease in H3K9me1, H4K5 and K12ac along with a reduction in total histone H3 and H4. HAT1 and RBBP7 were also decreased in response to PRPSAP1 depletion, without any noticeable changes in other histone chaperones, supporting the decrease in H4K5 and K12 acetylation upon PRPSAP1 depletion (Figure S4E).

Since we observed a direct interaction of PRPSAP1 with HAT1 and RBBP7, we speculated that PRPSAP1 might influence the acetyltransferase activity of HAT1-RBBP7. To test this, we performed an *in vitro* HAT assay using recombinant histone H4 as the substrate. The assay revealed that adding PRPSAP1 to the HAT1-RBBP7 complex enhances acetylation of histone H4 at both K5 and K12 residues (Figure 4L). HAT1 is known to regulate histone H4 levels^45^. The regulation of HAT1 activity by PRPSAP1 explains why depleting PRPSAP1 leads to reduced levels of H4 along with decreased acetylation at K5 and K12. Since acetylation at H4K5 and H4K12 is a critical step in the deposition of new histones, the regulation of HAT1-RBBP7 activity by PRPSAP1 underscores a crucial role for PRPSAP1 in this key step of histone deposition.

To dissect the distinct roles of PRPS1 and PRPSAP1 in the PRPS-PRPSAP complex and their involvement in histone regulation, we developed a dual differential degron system fusing PRPS1 to dTAG and PRPSAP1 to BromoTag at endogenous loci (Figure S5A). This system allowed us to selectively degrade PRPS1 or PRPSAP1 using dTAG47 or AGB1 within same cellular environment. We then asked if one component of the PRPS-PRPSAP complex affects the other’s interaction with histone chaperones. Upon depleting PRPSAP1 using AGB1, we observed a notably weakened interaction between PRPS1 and the histone chaperone NASP, both the tNASP and sNASP isoforms, even though PRPS1 can bind NASP directly *in vitro* without PRPSAP1 (Figure S5B). In contrast, depleting PRPS1 with dTAG47 did not affect PRPSAP1’s interactions with HAT1 (Figure S5B). Since PRPSAP1 can interact effectively with the HAT1-RBBP7 complex even in the absence of PRPS1, and because PRPSAP1 enhances the activity of HAT1-RBBP7 (Figure 4L), this explains why depleting PRPS1 but not PRPSAP1 leads to increased acetylation at H4K5 and H4K12. Taken together, these findings indicate that PRPSAP1 has a critical role in bridging the PRPS enzyme complex to histone chaperones.

### PRPS-PRPSAP complex regulates the engagement of histone chaperones with histones

Although we discovered a strong association of PRPS1 and PRPSAP1 with histone chaperones, we did not detect an interaction of PRPS or PRPSAP with histones themselves. We tested if PRPS1 governs the interaction of downstream histone chaperones with newly synthesized histones, as a potential mechanism for its effect on histone deposition. In PRPS1-dTAG HEK293T cells we expressed FLAG-tagged histone H3.3 or H4 under tetracycline induced promoter (Figure 5A). This system enabled us to induce new histone H3.3 or H4 expression in a controlled manner and selectively immunoprecipitate newly translated histones. The histone H3.1 and H3.3 variants utilize a common set of early histone chaperones, including NASP; therefore, we expect that the results obtained from experiments with H3.3 will also apply to H3.1. FLAG-tagged histone H3.3 or H4 were detectable as early as 4 hours post doxycycline treatment, indicating their status as newly synthesized histones (Figures 5B, C). We isolated these newly synthesized histone H3.3 or H4 through FLAG immunoprecipitation from the cytosolic fraction, adding an additional criterion to ensure their classification as new histones, from unperturbed or PRPS1 depleted cells. Our coimmunoprecipitation assays revealed a substantially reduced interaction of NASP, HAT1, RBBP7 and ASF1 with new histone H3.3 or H4 when PRPS1 was depleted (Figures 5D, E). Most importantly, interaction between H3.3 and H4, a crucial and early step in histone maturation, was also debilitated upon PRPS1 loss (Figures 5D, E).

**Figure 5.**
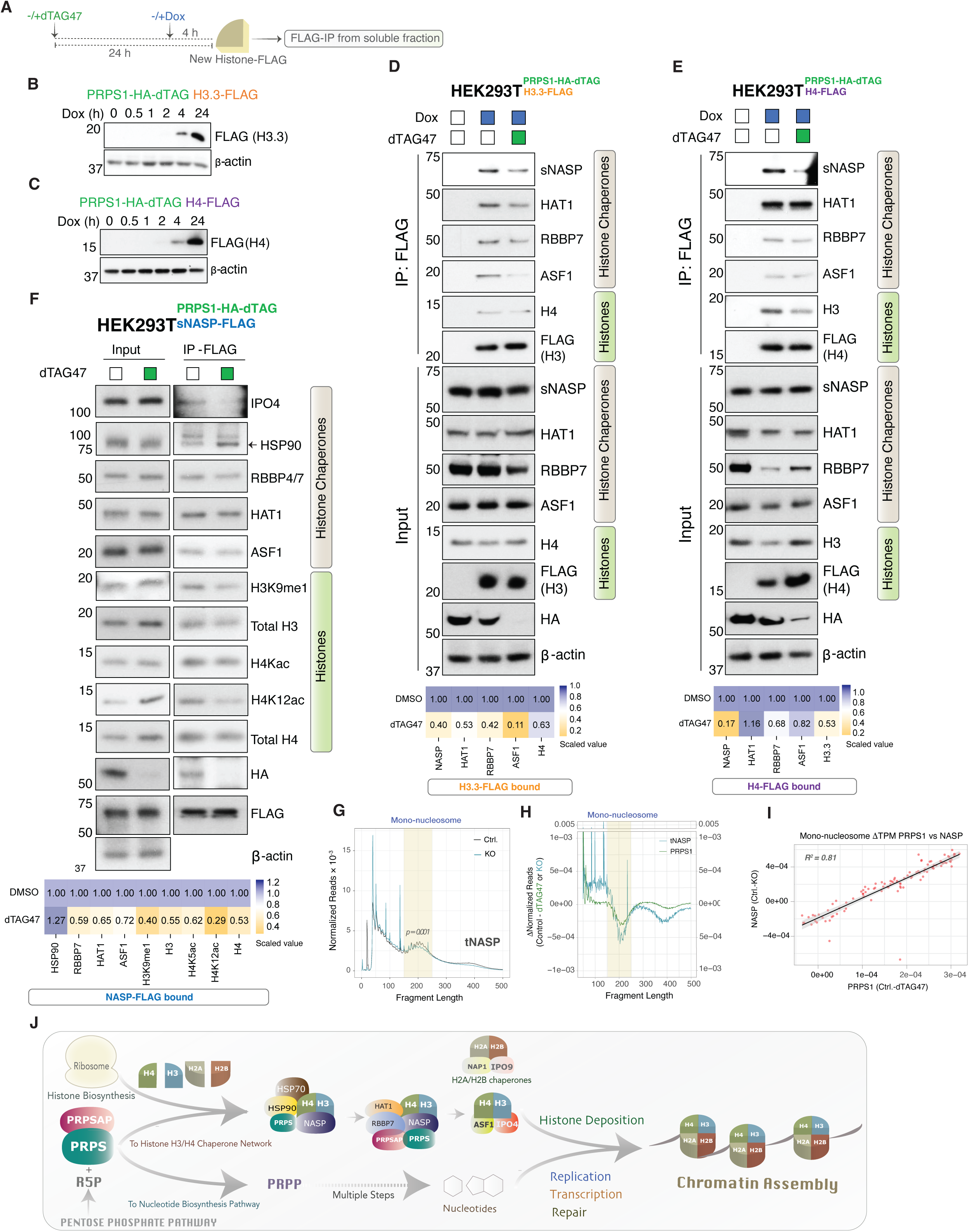
PRPS-PRPSAP complex regulates the engagement of histone chaperones with histones. **(A)** Schematic showing the expression of dox inducible histones and workflow to assess the interaction of new histones with histone chaperones. **(B-C)** Induction of FLAG-tagged H3.3 *(B)* and H4 *(C)* expression over time in response to doxycycline treatment. **(D-E)** Co-IP showing the interaction of new histone H3.3-FLAG *(D)* and H4-FLAG *(E)* with histone chaperones in soluble fraction from the cells pre-depleted with PRPS1 for 24 hours or in presence PRPS1. The heatmap below shows the relative levels of H3.3 or H4 bound histone chaperones with or without dTAG47 treatment, as quantified from the blots above. **(F)** Interaction of sNASP with histones and histone chaperones in HEK293T^PRPS1-HA-dTAG^ cells overexpressing FLAG-sNASP, with or without PRPS1 depletion for 24 hours. The heatmap below shows the relative levels of FLAG-sNASP bound histones and associated posttranslational makes and histone chaperones with or without dTAG47 treatment, as quantified from the blots above. **(G)** Fragment length distribution from nucleosome positioning analysis of ATAC-seq signal from WT or tNASP KO HEK293T cells^48^. Fragment lengths (150-250 bp) corresponding to mononucleosome is highlighted. *p* value was computed using Wilcoxon-singed rank test. **(H)** Comparison of changes in nucleosome positioning resulting from PRPS1 or tNASP loss analyzed by computing the difference in fragment length distribution of dTAG47 treated or tNASP KO samples to their respective controls. **(I)** Nucleosome positioning correlation with 95% confidence interval between PRPS1 depletion and tNASP KO analyzed by computing the difference in fragment length around mononucleosome (150-250 bps) of dTAG47 treated or tNASP KO samples to their respective controls. *Pierson correlation coefficient, R^2^ = 0.81.* **(J)** Model showing the coordinated regulation of nucleotide biosynthesis and histone deposition by PRPS-PRPSAP enzyme complex.

As a reciprocal approach, we expressed FLAG-tagged NASP in PRPS1-dTAG cells and conducted coimmunoprecipitation assays in presence or absence of PRPS1. A considerable reduction in NASP bound new histones as indicated by H3K9me1, H4K5 and K12ac were seen following PRPS1 depletion despite elevated levels of new histones in total lysate. In addition, the ability of NASP to interact with other chaperones including RBBP7, ASF1 and importin-4 was also diminished when PRPS1 was degraded (Figure 5F).

Histone chaperones play a crucial role in the histone deposition pathway by aiding in histone folding and protecting histones from degradation. If PRPS1 regulates the recruitment of histone chaperones to histones, then histones would be more susceptible to degradation under PRPS1-depleted conditions. To test this hypothesis, we performed an in vitro proteolysis assay using pronase, a non-specific protease^46^. We incubated newly synthesized histones H3 and H4, isolated from cells either depleted of PRPS1 or with intact PRPS, with varying concentration of pronase. New histones H3 and H4 purified from PRPS1-depleted cells were digested by pronase at lower concentrations compared to histones from PRPS1-intact cells (Figure S5C). This result suggests that in the absence of PRPS1, histones H3 and H4, which exhibit weakened interactions with histone chaperones, are more vulnerable to protease degradation, further emphasizing a critical role of PRPS1 in the histone chaperone pathway. Next, we assessed the impact of selective degradation of PRPS1 or PRPSAP1 on histone interactions with histone chaperones in HEK293T dual degron cells. Our findings revealed that the degradation of either PRPS1 or PRPSAP1 alone is sufficient to disrupt the association of both H3.3 and H4 with NASP (Figure S5D, E). These results are congruous with our observation that the absence of PRPSAP1 reduces the interaction between PRPS1 and NASP, supporting the idea that both PRPS and PRPSAP are essential for histone deposition.

Based on PRPS1 direct interaction with NASP and the defect in NASP engagement with histones upon PRPS1 depletion, NASP appears to be the main histone chaperone regulated by PRPS1. We investigated if NASP loss also results in nucleosome positioning defects, by conducting meta-analysis of tNASP ATAC-seq data^47^. Our analysis shows that tNASP loss leads to significantly reduced mono-nucleosomes, reflected by a reduced read count of fragment lengths between 150-250bp (Figure 5G). Notably, the reduction in mono-nucleosome read counts was more pronounced in NASP knockout cells than in PRPS1-depleted cells, as indicated by the greater difference in mono-nucleosome read counts in NASP knockout cells compared to PRPS1 depletion (Figure 5H). This more pronounced effect is likely due to NASP, as a histone chaperone, directly regulating histone deposition. Additionally, a constitutive NASP knockout cell line was used, while PRPS1 depletion was induced for 24 hours using the dTAG system. Nonetheless, the mono-nucleosome patterns resulting from PRPS1 or NASP loss were highly correlated (R² = 0.81) (Figure 5I), suggesting that PRPS1 loss phenocopies NASP knockout, both characterized by fewer mono-nucleosomes. These findings highlight the role for PRPS1 in nucleosome assembly through NASP.

Overall, our findings elucidate that PRPS-PRPSAP enzyme complex integrates into histone chaperone network and governs the recruitment of histone chaperones to histones and H3-H4 interaction, which are crucial early steps in histone deposition and chromatin assembly (Figure 5J).

## Discussion

The production of nucleotides for DNA replication, repair and transcription and the synthesis of histone proteins for the assembly of chromatin are major biosynthetic processes that are required for duplication of the genome and maintenance of genomic integrity. These critical processes have been presumed to function independently. However, our study identifies a novel and critical link between histone maturation and nucleotide metabolism through the PRPS-PRPSAP complex. Proper histone levels are critical for the maintenance of cell viability and protection against genomic instability. Previous work established PRPS enzymes as key players in nucleotide metabolism that catalyze the synthesis of PRPP from ribose-5-phosphate, the rate-limiting step in the nucleotide biosynthetic pathways^22^. Our work shows that the PRPS and PRPS-associated proteins (PRPSAPs), are integral members of the histone chaperone network, facilitating the proper folding, modification, and ultimately deposition of newly synthesized histones H3 and H4 into chromatin^22^. The loss of PRPS enzymes result in a marked accumulation of newly synthesized histone H3 and H4 marks in the cytosol and a corresponding decrease in their levels in the chromatin fraction. This indicates that PRPS enzymes are essential for early steps in histone maturation required for new histones into chromatin. iPOND experiments provided compelling evidence that function of PRPS proteins in the engagement of nascent histones with their chaperones leads to reduced deposition of histones onto replicating DNA, a process vital for maintaining chromatin integrity during DNA replication^8,11^. SNAP-tag experiments in asynchronous cells also show the accrual of nascent histones, including the replication independent H3.3 variant and H4 histones, and reduced their reduced incorporation into chromatin. These experiments suggest that PRPS proteins are required generally for nascent histone H3 and H4 availability outside of S-phase and will likely affect deposition of histones at sites of transcription, heterochromatin formation and DNA damage.

One of the most striking findings of our study is the functional independence of PRPS’s role in histone deposition from its catalytic activity in nucleotide biosynthesis. Our data showed that nucleotide biosynthesis pathway inhibition downstream to PRPS, does not impact histone marks, suggesting a distinct regulatory function of PRPS enzymes. This discovery expands the function of PRPS enzymes beyond its classical enzymatic role.

The maturation of histone H3 and H4 from translation to deposition into nucleosomes is a multistep process that requires a series of histone chaperone interactions^13,14^. Later steps in this process are specific to individual histone H3.1 and H3.3 variants, but the early steps are mediated by a common set of chaperones thought to contribute to histone folding^8,15,19^. Regulation at the early level provides broad control of histone availability for nucleosome assembly. Our study uncovers a previously unexplored function of PRPS enzymes by demonstrating that PRPS1, PRPS2, PRPSAP1 and PRPSAP2 are also critical components of the histone chaperone network that suggests that PRPS enzymes play a direct role in the early stages of histone processing, ensuring the proper folding and posttranslational modifications of nascent histones, critical for their stability and function.

Our data show that both PRPS and PRPSAP proteins contribute to nascent histone maturation However, the primary effects of PRPSAP depletion on histone levels differ from those observed with PRPS depletion, suggesting that PRPS and PRPSAP are essential for distinct aspects of the histone deposition pathway. Affinity purification and *in vitro* binding studies reveal distinct binding partners for PRPS and PRPSAP proteins within the histone chaperone network. PRPS1/2 are associated with NASP, whereas HAT1 and RBBP7 are associated with PRPSAP proteins. We show that PRPSAP1 enhances the lysine acetyltransferase activity of the HAT1-RBBP7 complex, specifically promoting acetylation of histone H4 at lysine residues K5 and K12. These modifications are crucial for nucleosome assembly, highlighting PRPSAP1’s essential role in coupling histone chaperone activity with chromatin formation. This highlights the multifaceted contribution of the PRPS-PRPSAP complex in chromatin assembly.

PRPS and PRPSAP proteins are known to form heteromeric complexes^48–50^. Loss of PRPS1 or PRPS2 results in an increased new H3 and H4, suggesting that the PRPS1-PRPS2 holoenzyme may be functionally required. Moreover, our data demonstrate that PRPSAP proteins are also required for assembly of the histone chaperone complexes. Therefore, we propose that the multimerized PRPS and PRPSAP proteins function as an *“assembleosome”* for the H3-H4-NASP-HAT1-RBBP7 histone chaperone complex, providing binding sites for distinct complexes required for the folding of the histone and modification of the histone tails designating them as new histones.

Since the seminal discovery linking cellular metabolism to histone acetylation^51^, and the exciting discovery linking IDH mutations to epigenetic reprogramming^52^, significant efforts have focused on exploring the connection between metabolism and chromatin. However, most studies have centered on the role of metabolite-driven post-translational modifications of histones in regulating chromatin functions like transcription. Our study shifts this paradigm by uncovering a direct link between chromatin assembly and nucleotide biosynthesis. PRPS and PRPSAP availability ensure that chromatin assembly is closely synchronized with DNA-dependent processes. When DNA synthesis begins, PRPS enzymes are activated to drive nucleotide production, and simultaneously, these enzymes play a central role in providing a substrate for histone-chaperone complex formation required for nucleosome deposition. This dual functionality provides a regulatory link between nucleotide biosynthesis and chromatin assembly, critical for maintaining genomic integrity. These findings significantly advance our understanding of the interplay between nucleotide metabolism and chromatin assembly, offering new perspectives on the regulation of these essential cellular processes.

### Limitations of the Study

Our study uncovers a critical mechanistic link between nucleotide and histone metabolism, both of which are required for chromatin assembly. While we observe a similar accumulation of newly synthesized histones upon PRPS1 or PRPS2 depletion, we do not distinguish whether this accumulation results from an overall reduction in PRPS enzyme levels (as PRPS1 and PRPS2 are highly similar and functionally redundant) or from disrupted PRPS1-PRPS2 complex formation.

Similarly, while we delineate distinct steps in the histone chaperone pathway regulated by PRPS1 and PRPSAP1, we have not yet defined the specific roles of PRPS1 and PRPS2 in this process.

We show that PRPS1 is enriched in NASP purification and demonstrate a robust interaction between PRPS1 and NASP both in vivo and in vitro. However, NASP was not identified in PRPS1 or PRPS2 affinity purifications, suggesting that only a small fraction of PRPS1 is bound to NASP.

Additionally, the fate of the accumulated histones observed in our study remains unknown. We primarily focused on the short-term and direct effects of PRPS or PRPSAP depletion on histone supply. While short-term depletion does not impact nucleotide biosynthesis, long-term depletion is expected to disrupt this pathway, which could have broader implications for histone metabolism. Future studies will be needed to explore these long-term effects.

## Resource Availability

All unique and stable reagents generated in this study will be made available on request by lead contact, Daniel R. Foltz with a completed materials transfer agreement.

## Data Availability

All data needed to assess the conclusions of this study are provided in the manuscript and supplementary materials. ATAC seq data will be available on GEO database upon publication.

## Acknowledgements

This work was supported by NIH grants GM143638 and U01CA260699. FACS was conducted with Ali Shilatifard lab and we thank Alex Lee for their assistance with FACS. Proteomics services were performed by the Northwestern Proteomics Core Facility, generously supported by NCI CCSG P30 CA060553 awarded to the Robert H Lurie Comprehensive Cancer Center, instrumentation award (S10OD025194) from NIH Office of the Director, and the National Resource for Translational and Developmental Proteomics supported by P41 GM108569. We thank Yuki Aoi from Shilatifard laboratory for helpful discussion on dTAG-degron *knock in*. We also thank Navdeep Chandel for their feedback on the manuscript.

## Authors Contribution

SS and DRF conceived the idea and planned the study. IBS provided the expertise in metabolism. VS provided the expertise in ATAC seq and DSC performed the analysis. SK performed the immunoprecipitation for NASP mass spectrometry, DRF acquired the confocal images and all the other experiments were conducted by SS. SS and DRF wrote the manuscript.

## Materials and Methods

### Cell Culture and Generation of dTAG *knock in* Cell Lines

HEK293T, HCT116 and DLD1 cells were grown in Dulbecco’s modified Eagle’s medium (DMEM) supplemented with 10% heat-inactivated fetal bovine serum (FBS)–optima (Atlanta Biologicals) and 1% penicillin/streptomycin (Gibco, Thermo Fisher Scientific). To generate a C-terminus FKBP12^F36V^ or BRD4BD^L387A^ knock-in, sgRNAs targeting PRPS, PRPS2 or PRPSAP1 were cloned into the Cas9-expressing vector pX330 (Addgene #42230) between BbsI sites. For the repair plasmid, we designed an epitope-tagged sequence (2x-HA for PRPS1 and 2x-V5 for PRPS2) fused to FKBP12^F36V^, followed by a cleaving peptide sequence (P2A) and an antibiotic resistance cassette (G418 for PRPS1 and blasticidin for PRPS2) to enable efficient identification of targeted cells. Similarly, for PRPSAP1-repair plasmid 2xV5 epitope tag was fused to BRD4BD^L387A^, followed by P2A sequence and blasticidin resistance cassette. This cassette was flanked by homology-directed repair arms, each approximately 500 base pairs in length: the left homology arm was located upstream of the stop codon, and the right homology arm started from the stop codon and extended immediately downstream. If required, a silent mutation was introduced in the PAM sequence to prevent Cas9 from cleaving the repair plasmid. The assembled sequence was cloned into the pTwist-Amp-high copy plasmid by Twist Biosciences for synthesis of the repair plasmid.

Target cells, plated in 6-well plates, were co-transfected with 500 ng sgRNA-Cas9 and 1500 ng repair plasmids using Lipofectamine™ 3000 transfection reagent (Invitrogen**-**L300000). After overnight transfection, fresh medium was replenished, and 24 hours later cells were plated in 10 cm plate. 24 hours post plating, antibiotic selection was started. Cells that survive antibiotic selection represent pooled population of hetero- and homozygous for knock in. To obtain cells with biallelic integration of epitope tag with FKBP12^F36V^ or BRD4BD^L387A^, antibiotic selected cells were plated in 15 cm plates with two very sparse dilutions; 125 and 250 cells and allowed to form colonies. Colonies were picked from trypsin disc when they were noticeable and transferred to 12-well plate. Each 12-well plate contain cells arising from single cell and hetero- and homozygous for knock in was validated through PCR using four independent primer-pairs covering the knock in junction and producing distinct product size. Note, both PRPS1 and PRPS2 are X-linked genes therefore, in principle it is not necessary to generate homozygous clones, when using the male cell lines. We used two male cell lines, HCT116 and DLD1 with PRPS1 or PRPS2-dTAG in our study and generated homozygous clones only for HCT116. Homozygous clones with PRPS or PRPS2-dTAG were generated for HEK293Ts, which is a female cell line. PRPSAP1 is found on chromosome 17 and to generate PRPSAP1-BRD4BD^L387A^, homozygous clones from single cell were generated for all the cell type used. To generate PRPS1-FKBP12^F36V^ and PRPSAP1-BRD4BD^L387A^ double knock-in, PRPSAP1-BRD4BD^L387A^ were introduced in HEK293T cells with PRPS1-FKBP12^F36V^ biallelic integration and homozygous clones were generated.

### Plasmids and Cloning

PRPS1 (TRCN0000355958) and PRPS2 (TRCN0000010130) shRNA were purchased from Millipore Sigma. C-terminus SNAP-FLAG tagged H3.3 and H4 were amplified using forward primer containing EcoRI site and reverse primer with FLAG-tag sequence and XbaI site and SNAP-3xHA tagged histone H3.3 and H4 in pBABE vector as templets, a gift from Lars Jansen. SNAP-FLAG tagged H3.3- and H4 PCR fragments were subsequently cloned into pLenti-III-PGK vector between EcoRI and XbaI sites. Doxycyclin inducible H3.3-3xFLAG and H4-3xFLAG were generated through Gateway cloning by introducing 3xFLAG-tag in pCW57.1 vector (Addgene#41939).

N-terminus FLAG-tagged PRPS1 or PRPS2 entry clones were generated by sub-cloning N-FLAG-PRPS1 (Sinobiologicals, HG17214-NF) and PRPS2 (Sinobiologicals, HG15128-NF) into pDONR221 vector (Invitrogen, 12536017). The entry clones were subsequently cloned into pLenti CMV Puro DEST (w118-1) (Addgene#17452) through Gateway cloning to generate lentiviral expression plasmid. The N-FLAG-PRPS1 was used as templet to generate ΔCD (aa 194-205) and ΔR5B (aa 213-228) mutant PRPS1 through sequential PCR. For ΔCD mutant, forward and reverse PCR primers were designed targeting the flanking region of catalytic domain, aa(187-193)+aa(206-211) skipping the catalytic domain. Two PCR fragments were generated using the forward primer covering the N-FLAG-PRPS1 (1-211) and PRPS1 (187-318), both lacking the catalytic domain (194-205). These two PCR fragments were further used as templet in 1:1 ratio to generate ΔCD mutant using the primer pair targeting N and C-terminus of PRPS1. ΔR5P mutant was similarly generated using the primer pair targeting the flanking region of ribose-5-phosphate domain aa(206-212)+(229-234), skipping the ribose-5-phosphate domain aa(213-228). PRPSAP1 (isoform 1) entry clone (HsCD00288768) and PRPSAP2 (HsCD00515561) from were purchased from DNASU, and FLAG-PRPSAP1 or PRPSAP2 expression clone was generated through LR reaction with pLenti6.2-3xFLAG-V5 (Addgene, 87072). Generation of 3x-FLAG-sNASP and 3x-FLAG-tNASP are described earlier ^53^.

### Lentivirus Production and Stable Cell Line Generation

For lentivirus production, desired transfer plasmid (1 μg) was cotransfected with 0.5 μg of VSV-G coat protein vector, pMD2.G (Addgene, 12259), and 1 μg of psPAX2 (Addgene, 12260) in HEK293T cells plated in 12-well plate using Lipofectamine 3000 (Thermo Fisher Scientific, L3000008) reagent. After overnight transfection, fresh medium was replenished, and 24 hours later, the first batch of viral supernatant was collected and stored in 4°C. Cells were replenished with fresh medium again, and the final batch of lentivirus supernatant was collected 24 hours later. Both the viral batches were pooled and filtered through 0.45-μm surfactant-free cellulose acetate membrane filter. As needed, lentiviral particles were either stored in −80°C or used immediately for transduction. To generate HEK293T cells stably expressing FLAG-tagged FLAG-PRPS WT, ΔCD, ΔR5P mutants, PRPS2, PRPSAP1, PRPSAP2, s- or tNASP, HCT116 cells expressing FLAG-SNAP tagged histone H3.3 or H4, HEK293T^PRPS1-HA-dTAG^ or HEK293T^PRPS1-HA-dTAG^ ^PRPSAP1-V5-BromoTag^ cells expressing doxycycline inducible FLAG tagged H3.3 or H4, 100 × 10^3^ cells were seeded a day before transduction. Cells were incubated with lentivirus with polybrene (8 μg/ml) overnight. Cells were then replenished with fresh medium and grown for 24 hours followed by appropriate antibiotic selection.

### Immunoprecipitation and Western blot

FLAG immunoprecipitation, elution of FLAG-immunoprecipitate by FLAG peptide and Western blots were performed as described earlier^29^. For immunoprecipitation of PRPS1-HA-dTAG and PRPSAP1-V5-BromoTag, anti-HA (Pierce 88836) and anti-V5 (Proteintech v5tma) magnetic beads were used and immunoprecipitates were boiled in 4x-Laemmli sample buffer before analyzing through SDS-PAGE. Remaining steps in immunoprecipitation were same as FLAG-immunoprecipitation as described earlier. To purify new FLAG-tagged histone H3.3 or H4 from HEK293T^PRPS1-HA-dTAG^ or HEK293T^PRPS1-HA-dTAG PRPSAP1-V5-BromoTag^, cells were pre-treated with dTAG47, AGB1 or both for 20 hours and then supplemented with 2 μM doxycycline for 4 hours. New histones were FLAG immunoprecipitated from the prenucleosomal fraction, eluted with FLAG peptide and interaction with histone chaperones were examined by immunoblot. For Pronase proteolysis assay, FLAG-tagged histone H3.3 or H4 purified from the prenucleosomal fraction were treated with varying doses of Pronase (Roche 10165913103) as mentioned in the Figure for 45 minutes and histone proteolysis was examined through immunoblotting with anti-FLAG antibodies.

### Isolation of Newly Translated Proteins

HEK293T^PRPS1-HA-dTAG^ cells with or without dTAG47 treatment for 21 hours, were depleted from methionine using methionine free media (Thermo Fisher Scientific, 21013024) for 1 hour and subsequently supplemented with L-azidohomoalanine (Thermo Fisher Scientific, C10102) for 2 hours. For PRPS1-depleted samples, dTAG47 was maintained throughout with methionine depletion and AHA treatment, and the combined timeframe of all the treatments was 24 hours. Cells were harvested and click reaction was performed following using Click-iT^TM^ protein reaction buffer kit (Thermo Fisher Scientific, C10276) following manufacturer’s protocol. Lysates after click reaction were subjected to immunoprecipitation with streptavidin C1 dynabeads (Invitrogen 65001) and analyzed through immunoblotting with antibodies as indicated.

### Antibodies

PRPS1 (Thermo Scientific 15549-1-AP and Proteintech, 15549-1), PRPS2 (Thermo Scientific PA5-42007 and Proteintech, 27024-1), PRPSAP1 (Proteintech 16790-1-AP), Histone H3 (Cell Signaling Technologies, 4499S), H3K9me1 (Abcam ab8896), Histone H4 (EMD Millipore 05-858), H4K5ac (RevMAb Biosciences 31-1081-00), H5K12ac (Cell Signaling Technologies13944S), Cyclin E2 (Cell Signaling Technologies 4132), Cyclin A2 (Cell Signaling Technologies 4656), α-tubulin (Proteintech, 66031-1), β-actin (Sigma-Aldrich, A2228), FLAG (Sigma-Aldrich, F3165), HÁ (Cell Signaling Technologies 2999), V5 (Thermo Fisher Scientific R961-25), Streptavidin (Jackson ImmunoResearch 016 030-084), SNAP (New England Biolabs P9310S), NASP (Abcam ab181169), HSP90 (Cell Signaling Technologies 4874), Importin4 (Proteintech 11679-1-AP), HAT1 (Cell Signaling Technologies 15348S and Proteintech 11432-1-AP), RBBP7 (Cell Signaling Technologies 4522), RBBP4/7 (Cell Signaling Technologies 9067S), ASF1 (Cell Signaling Technologies 2990S), GST (Proteintech, HRP 66001), 6xHis (Proteintech HRP66005).

### Quantitative Reverse Transcription PCR

HEK293T^PRPS1-HA-dTAG^ cells treated with DMSO or dTAG47 for 24 hours. Total RNA was isolated using an RNA extraction kit (Qiagen, 74106), and complementary DNA (cDNA) was prepared form 1 μg of RNA using an iScript cDNA synthesis kit (Bio-Rad, 1708890). cDNA was diluted 10 times, mixed with iTaq universal SYBR green supermix (Bio-Rad, 1725121), and subjected to real-time quantitative PCR using standard procedure using histone gene specific primers. Gene expression analysis was performed using 2^−ΔΔ*CT*^ method.

### Cell Cycle Analysis

Control of dTAG47 treated cells were fixed 70% ethanol in 1x PBS and stored in 4^0^C overnight. Ethanol was removed and cells were wash with 1x PBS. Cells were then stained with propidium iodide solution containing 0.01 mg/ml propidium iodide and 250 mg/ml RNaseA and DNA content were measured through flow cytometry.

### Subcellular Fractionation

Subcellular fractionation to isolate cytosolic and chromatin bound proteins were performed as described by Gillotin et al.^54^. Briefly, cell pallets were collected by scrapping the cells in ice cold PBS. Cell pallet were lysed in 5 cell pallet volume of buffer E1(50 mM HEPES-KOH; pH 7.5, 140 mM NaCl, 1 mM EDTA; pH 8.0, 10% glycerol, 0.5% NP-40, 0.25% Triton X-100, 1 mM DTT supplemented with 1x protease inhibitor cocktail), centrifuged at 1,100 *× g* at 4 °C for 2 min and supernatant were collected in a fresh tube representing cytoplasm (soluble or prenucleosomal) fraction. Remaining pallet was washed twice in E1 buffer at 1,100 *× g* at 4 °C for 2 min with 10 minutes incubation in E1 buffer on ice before the second wash. After discarding the supernatant, pallet was dissolved in same volume of E2 buffer (10 mM Tris-HCl; pH 8.0, 200 mM NaCl, 1 mM EDTA; pH 8.0), 0.5 mM EGTA; pH 8.0, supplemented with 1x protease inhibitor cocktail) and centrifuged at 1,100 *× g* at 4 °C for 2 min. The supernatant containing the nuclear soluble proteins were not preserved in our study and discarded. The remaining pallet were washed twice with E2 buffer same as the pallet washed with E1 buffer in the previous step. The pallet was resuspended in 2.5 cell pallet volume of E3-benzonase buffer (50 mM Tris-HCl; pH 7.5, 20 mM NaCl, 1 mM MgCl_2_, 1% NP-40, supplemented with 1x protease inhibitor cocktail and benzonase in 1:1000 dilution when ready to use and incubated on rotating platform for 20 minutes to liberate chromatin bound proteins. Finally, both cytosolic and chromatin fractions were centrifuged at maximum speed at 4 °C for 10 minutes and supernatants were transferred to fresh tubes.

### Isolation of Proteins on Nascent DNA (iPOND)

Native iPOND was performed as described by Sirbu et al^43^. Briefly, HEK293T, HCT116 and DLD1 cells with PRPS1-HA-dTAG knock in were grown in 10 cm dishes and treated with 500 nM dTAG47 or left untreated. 23 hours 20 minutes post dTAG47 treatment, cells were labeled with 10 μM EdU for 10 min followed by 10 μM thymidine chase for 30 min bringing dTAG47 treatment for 24 hours, before cells were harvested. A small fraction of cells collected from the same dish to harvest total protein to examine PRPS1 depletion, while remaining cells were subjected to click reaction as previously described using biotin azide. No click reaction was included as negative control. After micrococcal nuclease digestion chromatin was extracted and subjected to streptavidin immunoprecipitation using streptavidin C1 dynabeads (Invitrogen 65001) overnight on rotator in cold room. Beads were washed five times with extraction buffer and bead bound proteins were released by addition of equal volume of 2x Laemmli sample buffer followed by heating at 95°C for 10 mins. Recovered proteins were subjected to SDS-PAGE and immunoblotted for histones.

### SNAP Labelling and Microscopy

SNAP labeling was performed as described previously^41^. HCT116^PRPS1-HA-dTAG^ cells stably expressing FLAG-SNAP-tagged histone H3.3 or H4 were grown in chambered cell culture slide (MatTek Corporation CCS-8) and treated with or without dTAG47 for 24 hours to deplete PRPS1. Cells were then blocked by 10 µM SNAP-Cell Block (New England Biolabs S9106S) for 30 minutes to quench pre-existing SNAP-tagged histones. Cells were washed twice with 1x-PBS and replenished with fresh media to allow the synthesis of new SNAP-tagged histones and chased for 2 hours. 2 µM SNAP-Cell TMR Star (New England Biolabs S9105S) was added to cells for 30 minutes to label new SNAP-tagged histones. Cells were washed twice with 1x-PBS and straight fixed in 4% paraformaldehyde for 10 minutes at room temperature or pre-extracted with 0.2% triton X-100 in cytoskeletal buffer for 3 minutes before fixation. Fixed cells were washed twice with 1x-PBS, mounted on VECTASHIELD antifade mounting media with DAPI (Vector Laboratories H-1800-2) and imaged on Zeiss LSM800 confocal microscope.

Confocal and Airyscan images were collected on a Zeiss LSM800 confocal Microscope using a 63x oil-immersion objective lens. Images were processed as orthogonally projected stacks of maximum pixel values using the Zeiss Zen software. Channels were scaled identically within panels and conditions of Figures. Quantification of whole-cell and nuclear intensity was conducted using ImageJ. Nuclear intensity in pre-extracted cells was determined by identifying nuclear volume based on DAPI staining and measuring the integrated intensity of the TMR-Star signal within that nuclear region. To determine the total non-chromatin and chromatin bound pool of new histone, the TMR-Star signal was determined from straight-fixed cells. Total integrated TMR-Star intensity was measured across the entire cell. *p* values were computed using Student’s t-test.

### HAT Assay

HAT assay was carried out in HAT buffer containing 50 mM Tris HCl pH 8.0, 50 mM KCl, 5% glycerol, 0.1 mM EDTA, 1 mM DTT, 10 mM sodium butyrate, 200 μM acetyl coenzyme-A (Millipore Sigma A2056) and 1x protease inhibitor cocktail HAT1 as catalytic enzyme, RBBP7 and PRPSAP1 as cofactors and recombinant histone H4 a substrate. Briefly, 250 ng recombinant histone H4 (EpiCypher 15-0304) was incubated in 100 ng 6xHis-HAT1, aa 20-341 (Novus Biologicals, NBP1-41234), 325 ng 6xHis-SUMO-RBBP7 (CUSABIO Technology, CSB-EP621959HU) and 6xHis-PRPSAP1 (Proteintech, Ag10238) in 25µl KAT buffer at room temperature for 1 hour. The reaction of stopped by the addition of 4x-Laemmli sample buffer and ¼ of the reaction subjected to SDS-PAGE. HAT1, RBBP7, PRPSAP1 and H4K5 and K12ac acetylation was detected through immunoblotting using specific antibodies. Memcode staining (Thermo Scientific 24585) was performed to assess the total histone H4 used in the assay.

### GST Pulldown Assay

1 μg of GST (Millipore Sigma SRP5348), GST-NASP (Abnova, H00004678-P01), or GST-PRPSAP1 (Proteintech Ag10227) was incubated to GST magnetic beads (Thermo Scientific, 78601) for 1 hour on rotator. Non-covalently GST proteins bound to glutathione beads were washed three times with NETN buffer (20 mM Tris-HCl pH 7.5, 150 mM NaCl, 0.5% Nonidet P-40, and 1x-protease inhibitor cocktail). Bead bound GST or GST-NASP was incubated with 250 ng 6xHis-PRPS1(Novus Biologicals, NBP1-37079) overnight in cold room with rotation. Similarly, bead bound GST or GST-PRPSAP1 (Proteintech, Ag10227) was incubated with 250 ng 6xHis-HAT1, aa 20-341 (Novus Biologicals, NBP1-41234) or 6xHis-SUMO-RBBP7 (CUSABIO, CSB-EP621959HU). Next day, beads were washed five times with NETN buffer and binding were examined by immunoblot. Note, HAT1 interaction was detected using anti-HAT1 (Proteintech, 11432-1-AP) that is raised against HAT1 peptide (aa 241-419). Other antibodies used in GST pulldown assay are described in *‘Antibodies’* section.

### ATAC-seq

ATAC-seq on HEK293T^PRPS1-HA-dTAG^ cells treated with DMSO or dTAG47 for 24 hours was conducted as described previously^55^ and analyzed using nf-core/atacseq pipeline version 2.1.2^56^. In this pipeline, adapters were trimmed using trimgalore and reads were aligned to hg38 using BWA (v.0.7.17). HEK293T WT and tNASP KO samples ^47^ were mapped with BWA (v.0.7.17) with the parameters ’bwa mem -t 8’ to GRCh37 hg19 (UCSC). PRPS and NASP bam files were sorted with samtools (1.10.1) ’sort’. Duplicated reads were then marked and removed with picard-tools-2.21.4 Mark Duplicates (https://broadinstitute.github.io/picard/) and fragment length was calculated using the function fragSizeDist from ATACseqQC60 (v.1.26.0). The normalized values were calculated by dividing the observed fragment count for each fragment length by the total number of fragments using the function fragSizeDist2.

### FLAG-affinity Purification and Mass Spectrometry

FLAG-affinity purification followed by TCA precipitation was conducted as described previously^29^. Dried TCA precipitated pellets were resuspended in loading buffer and incubated at 95 °C for 10 minutes with shaking. Each sample was loaded into SDS-PAGE gel well and electrophoresed approximately 1 cm to form a concentrated gel plug. Gel plugs were excised and washed with high purity water followed by an in-gel digestion method where proteins were reduced (10 mM dithiothreitol), alkylated (20 mM iodoacetamide), and alternatively washed with 100 mM ammonium bicarbonate and acetonitrile. Proteins were digested overnight with 0.5 µg of trypsin at 37 °C, followed by acidification with formic acid to halt the digestion. The resulting peptide solution was dried by vacuum centrifugation. Dried peptides were resuspended in 30 µl of 0.1 % aqueous formic acid, shaken and sonicated for 10 minutes each, and centrifuged for 5 minutes at 20,000 ×g. 25 µl of peptide supernatant was transferred to a glass auto-sampler vial then placed in the auto-sampler of a Thermo Scientific Ultimate 3000 nano liquid chromatography instrument. 4 µl of sample was loaded onto an in-house packed C18 trap column using a 2.5 µl/min flow rate (0.1 % aqueous trifluoroacetic acid) then transferred to a C18 analytical column (75 µm × 25 cm). Peptides were separated using an increasing gradient of organic mobile phase (80 % acetonitrile and 0.1 % aqueous formic acid) and nano-electrosprayed into the heated source of a Thermo Scientific Orbitrap Elite mass spectrometer. The 15 most abundant peptides per survey (MS1) scan were isolated and fragmented (MS/MS) over a 120-minute method. Raw data files were converted to .mgf files and searched against the human proteome database in Mascot 2.7 software. Search results were imported into Scaffold 5 software.

To analyze the MS data, 1 was added to all spectral counts to ensure there were no zero values. For each condition, the total spectral count was calculated by summing all peptide spectral counts, and each individual spectral count was normalized by dividing it by this total. The mean normalized spectral count was then calculated from three replicates. The mean log2 fold change (FLAG/Parental) was determined across the three replicates. *p*-values were calculated using an unpaired t-test with Welch’s correction. Finally, a volcano plot was created by plotting the log_2_(fold change) on the x-axis and -log_10_(*p*-value) on the y-axis.

### Epiproteomic Histone Modification Panel

1.0 × 10^6^ HEK293T cells, transduced with non-targeting, PRPS1 or PRPS2 shRNA, were harvested in 1x PBS. Cell pellets were lysed using 0.3 % NP-40 then centrifuged to pellet the nuclei. Histones were extracted from the nuclei by addition of 200 mM sulfuric acid followed by centrifugation at 4,000 ×g for 4 minutes. The supernatant was transferred to a new tube and histone precipitation was performed using 20 % trichloroacetic acid on ice for 1 hour. Histone pellets were collected by centrifugation at 10,000 ×g for 5 minutes at 4 °C, washed with cold acetone and 0.1 % hydrochloric acid, and allowed to air dry. Purified histone pellets were resuspended in 100 mM ammonium bicarbonate. Unmodified lysine residues and protein N-termini were chemically derivatized with a solution of propionic anhydride in propanol for 1 hour at 51 °C. Samples were dried by vacuum centrifugation then resuspended in 100 mM ammonium bicarbonate. Histones were digested overnight with 0.5 µg of trypsin at 37 °C and the resulting peptide solution was dried by vacuum centrifugation. Dried peptides were resuspended in 100 mM ammonium bicarbonate, subjected again to chemical derivatization to propionylate newly exposed N-termini of peptides, then dried by vacuum centrifugation. Dried histones were resuspended in 50 µl of 0.1 % aqueous trifluoroacetic acid, shaken for 5 minutes, and centrifuged for 5 minutes at 20,000 ×g. The peptide supernatant was transferred to a glass auto-sampler vial then placed in the auto-sampler of a Thermo Scientific Ultimate 3000 nano liquid chromatography instrument. 2 µl of sample was loaded onto an in-house packed C18 trap column using a 2.5 µl/min flow rate (0.1 % aqueous trifluoroacetic acid) then transferred to a New Objective Picochip C18 analytical column (75 µm × 10.5 cm). Peptides were separated using an increasing gradient of organic mobile phase (80 % acetonitrile and 0.1 % aqueous formic acid) and nano-electrosprayed into the heated source of a Thermo Scientific Altis triple quadrupole Elite mass spectrometer. Technical triplicates were acquired, and the acquisition sequence was randomized. Data were collected using a multiple-reaction monitoring (MRM) targeted approach for an in-house developed list of precursor and product ions over a 60-minute method. Thermo raw data files imported in Skyline software and analyzed with an additional in-house developed software (Protein Assay). To analyze Epiproteomic data, mean peptide abundance from three replicates of control, PRPS1 or PRPS2 shRNA were calculate. PRPS1 and PRPS2 shRNA mean peptide abundance were normalized to control shRNA and heatmap was generated using ggplot2 package in R.

### Oligoes used in cloning

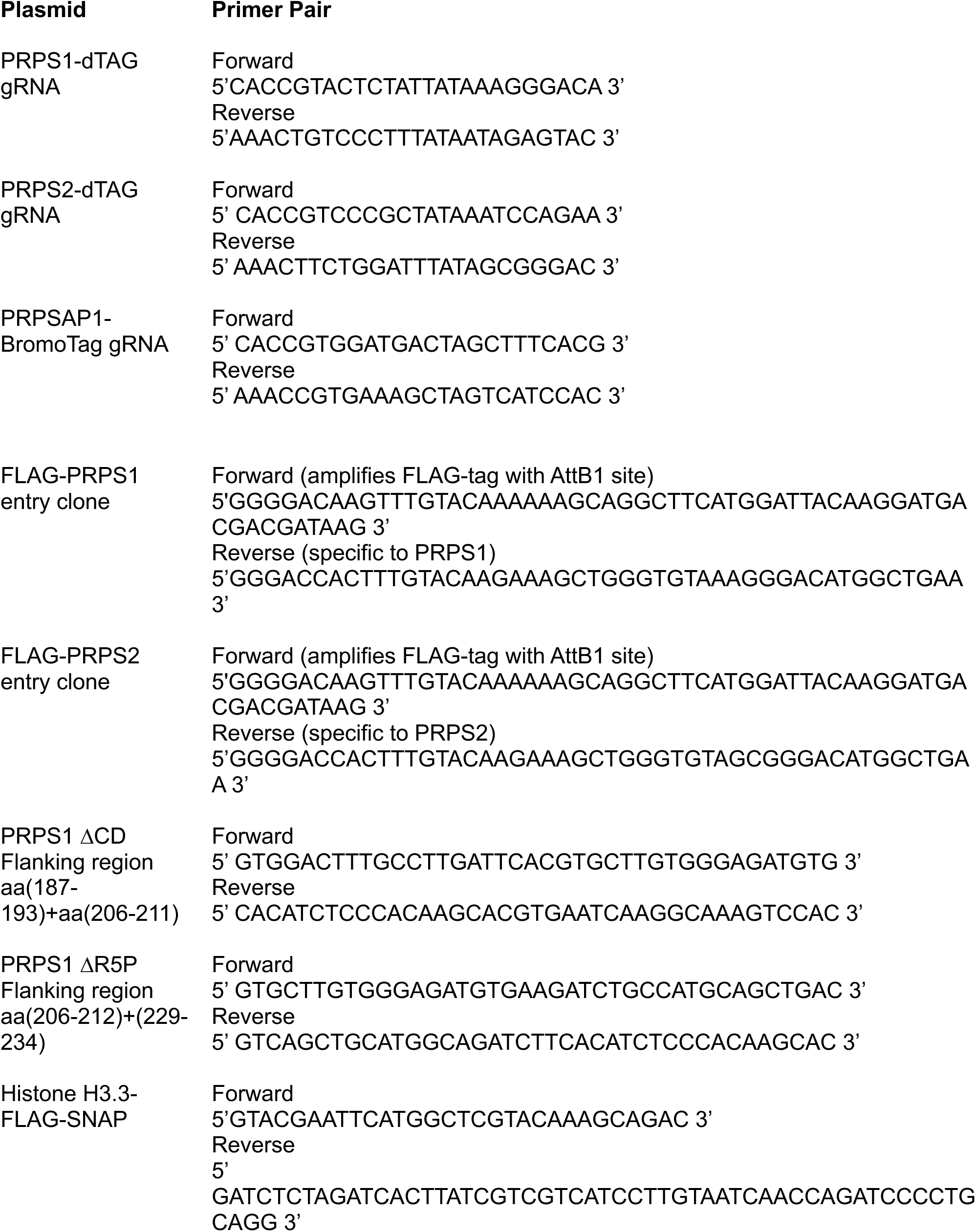
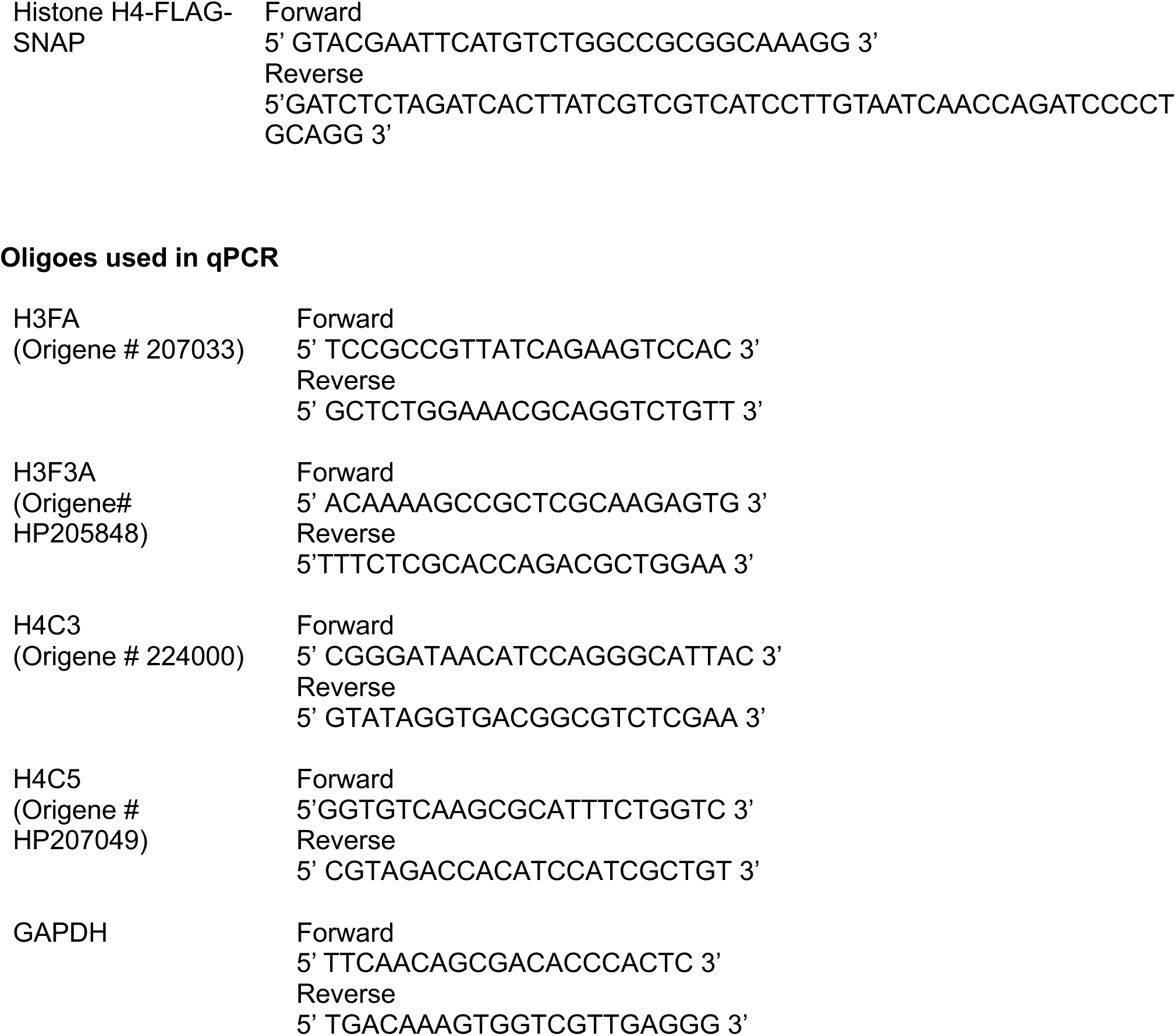

