## Supplemental Data for "Rate Limiting Enzymes in Nucleotide Metabolism Synchronize Nucleotide Biosynthesis and Chromatin Formation"

#### Supplementary Figure S1

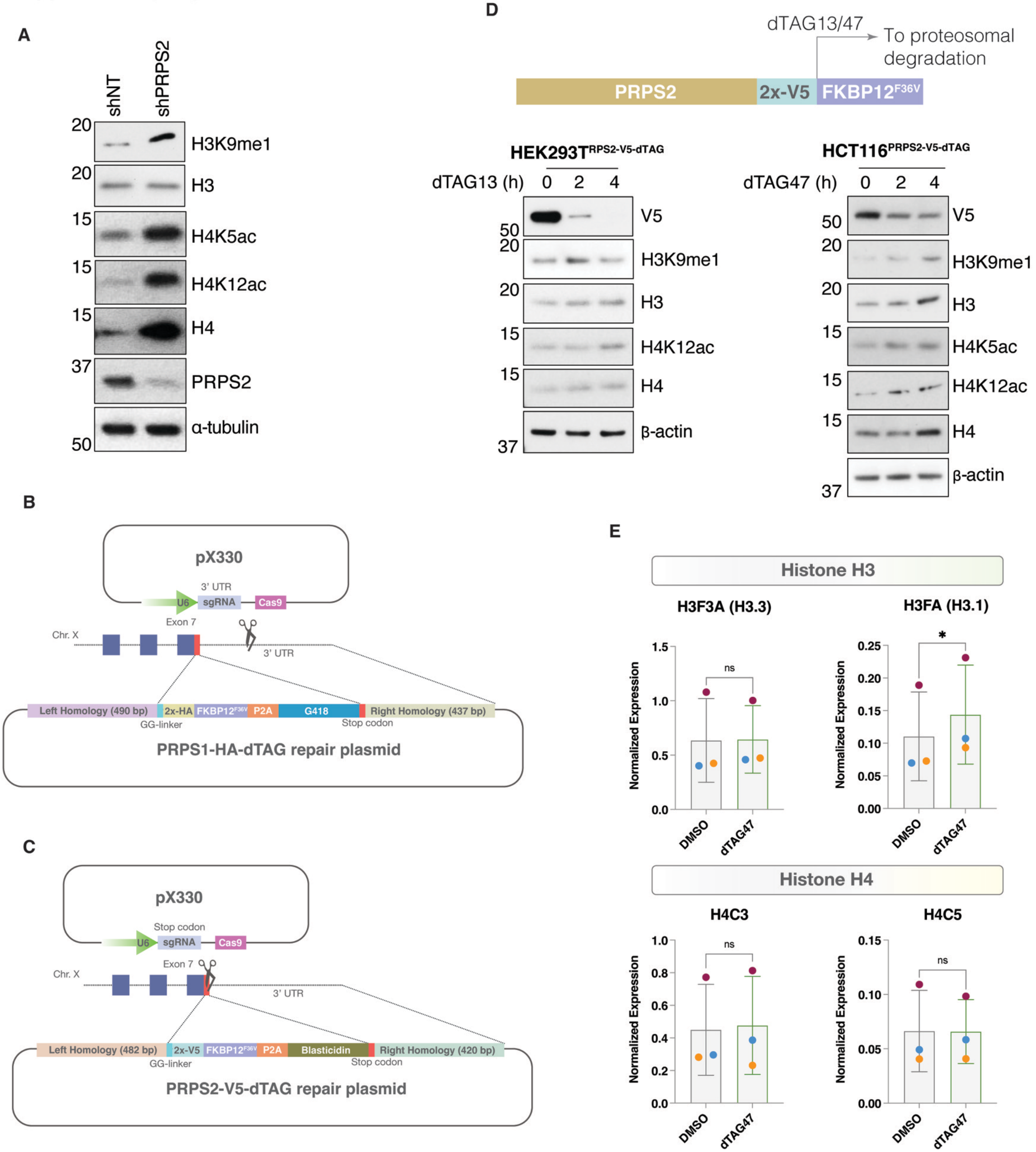

**Figure S1. shRNA mediated or dTAG-degron mediated PRPS loss result in increased total histone H3 and H4 and, new H3 and H4 marks. (A)** Total histone and posttranslational modification marks associated with new histones in Ctrl, PRPS2 shRNA HEK293T cells. **(B-C)** Strategy depicting PRPS1-HA- $FKBP12^{F36V}$ *(B)* or PRPS2-V5-$FKBP12^{F36V}$ *(C)* knock in through homology directed repair. **(D)** Schematic and induction of PRPS2-V5-dTAG proteasomal degradation in response to dTAG13/47 *(top)*. Total histone and posttranslational modification marks associated with new histones in response to time dependent depletion of PPRS2 in $HEK293T^{PRPS2\text{-}V5\text{-}dTAG}$ and $HCT116^{PRPS2\text{-}V5\text{-}dTAG}$ cells. **(E)** qPCR showing the mRNA levels of two of each histone H3 and H4 genes in $HEK293T^{PRPS1\text{-}HA\text{-}dTAG}$ cells with or without PRPS1 depletion mediated by dTAG47 treatment for 24 hours.

#### Supplementary Figure S2

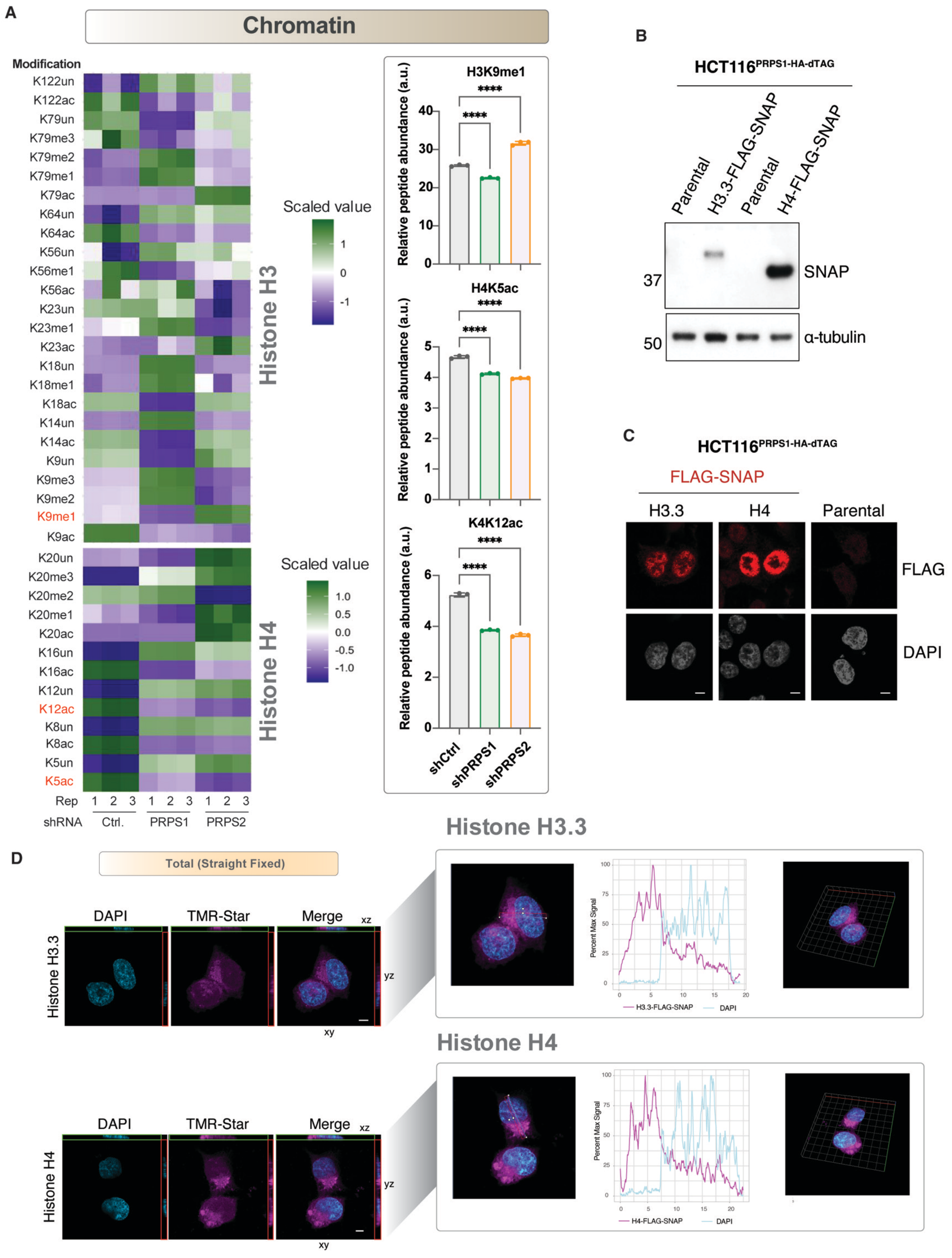

**Figure S2. PRPS loss reduces new histone marks and impacts new histone incorporation in chromatin. (A)** Epiproteomic Histone Modification Panel showing changes in histone modification changes in chromatin fraction in HEK293T cells transduced with ctrl., PRPS1 or PRPS2 shRNA. New histone marks H3K9me1, H4K5ac and H4K12ac are highlighted and changes in these marks are also shown through bar plot alongside. *n=3*, *p* values were computed using Student's t test, *** $p \leq 0.001$ **(B)** Expression of histone H3.3-FLAG-SNAP or H4-FLAG-SNAP in HCT116$^{PRPS1\text{-}HA\text{-}dTAG}$ background as detected though immunoblotting with anti-SNAP antibodies.**(C)** Immunofluorescence showing the distribution of histone H3.3-FLAG-SNAP and H4-FLAG-SNAP in HCT116$^{PRPS1\text{-}HA\text{-}dTAG}$ background as stained with anti-FLAG antibodies. *Scale bar: 5 μm* **(D-E)** Orthogonal projection of straight fixed histone H3.3-FLAG-SNAP *(D)* and H4-FLAG-SNAP *(E)* from *Figure 2D* showing non-overlapping TMR-Star and DAPI signal. Pseudo-colors were used to enhance the contrast, *scale bar: 5 μm.*

### Supplementary Figure S3

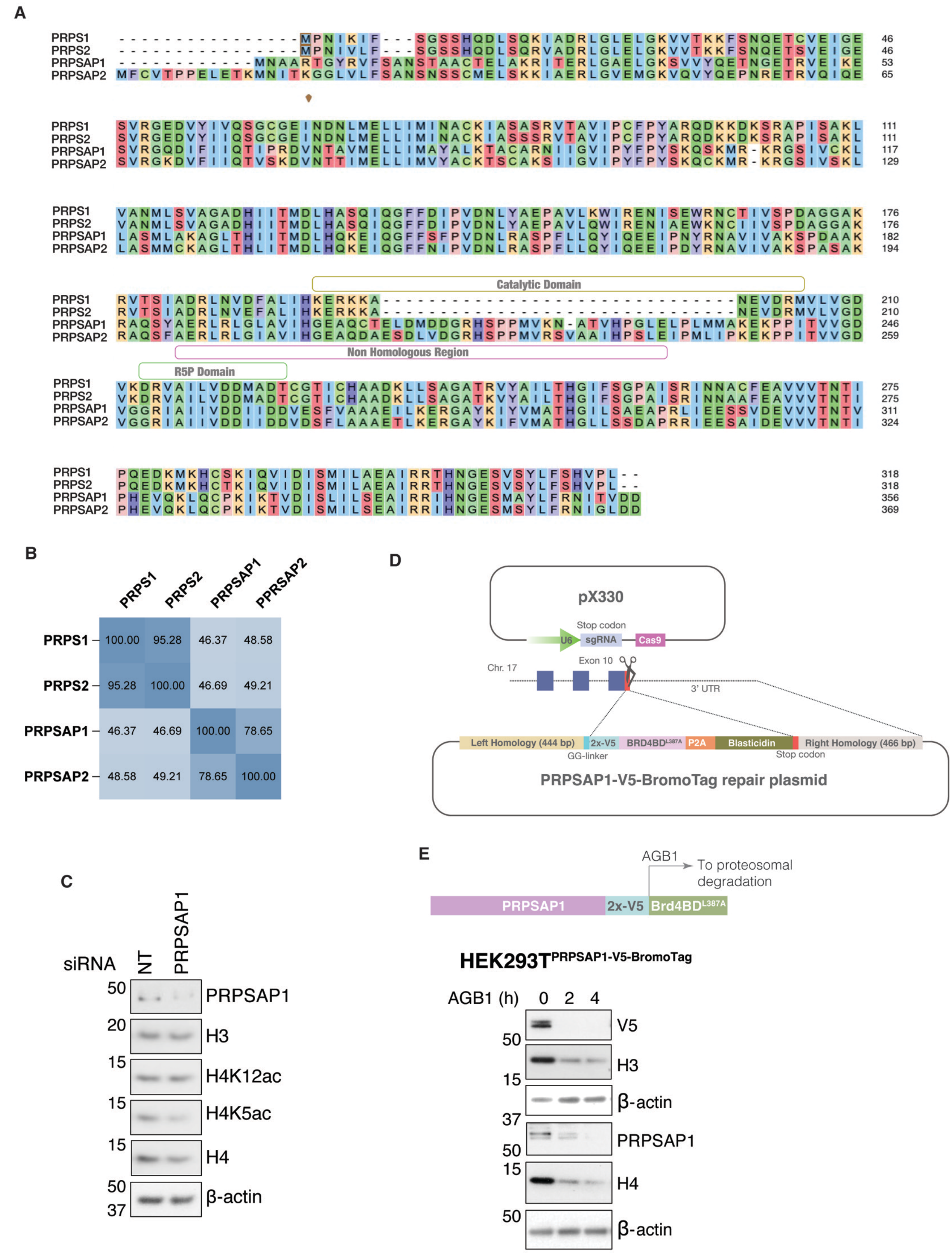

**Figure S3. PRPSAP1, as a component of PRPP synthetase, is involved in histone regulation. (A)** Sequence alignment of human PRPS1, PRPS2, PRPSAP1 and PRPSAP2 proteins. **(B)** Matrix showing the amino acid identities among PRPP-synthetase complex proteins. **(C)** Total histone and posttranslational modification marks associated with new histones in HEK293T cells transfected with non-targeting (NT) or PRPSAP1 siRNA for 48 hours. **(D)** Strategy depicting PRPSAP1-V5-BRD4BD$^{L387A}$ (BromoTag) knock in through homology directed repair. **(E)** Schematic and induction of PRPSAP1-V5-BromoTag proteasomal degradation in response to AGB1 *(top)*. Total histone H3 and H4 in response to time dependent depletion of PPRSAP1 in HEK293T$^{PRPSAP1\text{-}V5\text{-}BromoTag}$.

#### Supplementary Figure S4

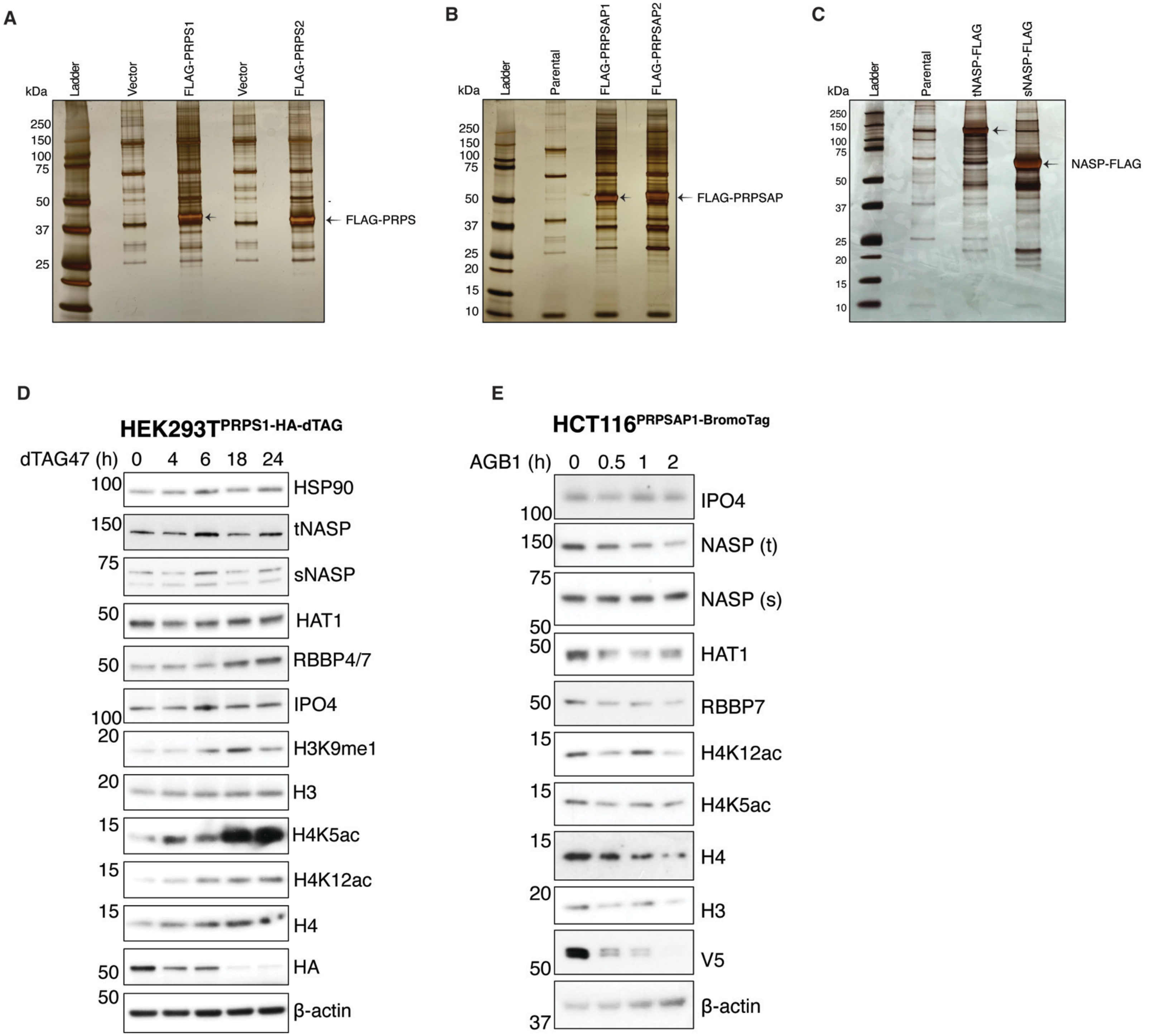

**Figure S4. PRPS-PRPSAP complex interacts with and regulates histone chaperones. (A-C)** Silver stain showing the FLAG-affinity purification of PRPS1 and PRPS2 *(A)* PRPSAP1 and PRPSAP2 *(B)* and tNASP and sNASP *(C)* from HEK293T cells. **(D)** Histones and histone chaperones levels as in response to PRPS1 depletion over time from HEK293T cells with PRPS1-HA-dTAG knock in. **(E)** Histones and histone chaperones levels as in response to PRPSAP1 depletion over time from HCT116 cells with PRPSAP1-V5-BromoTag *knock in*.

#### Supplementary Figure S5

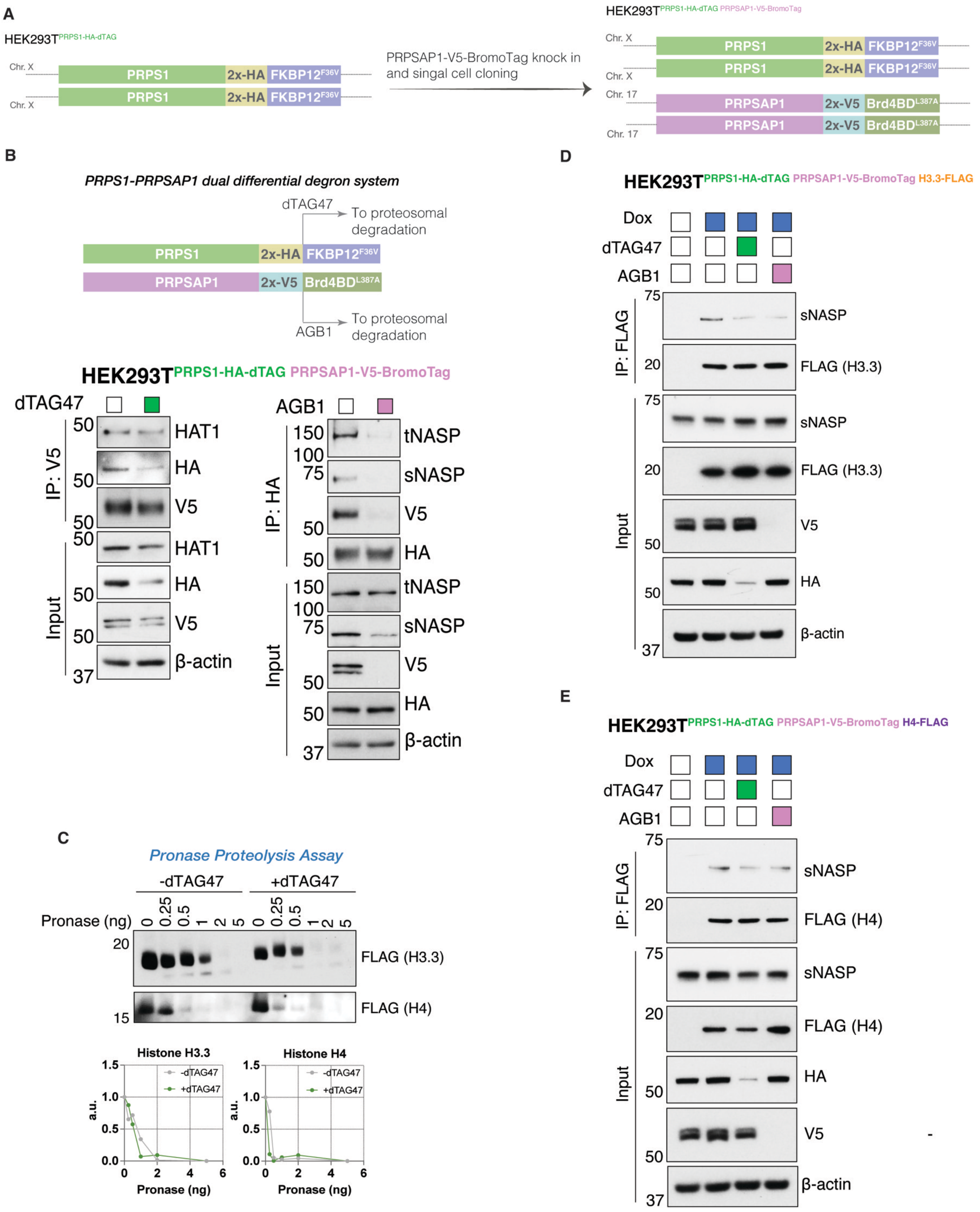

**Figure S5. Selective degradation of PRPS1 or PRPSAP1 to assess PRPP-synthetase complex role in histone regulation. (A)** Schematic showing the strategy for generation of PRPS1-HA-dTAG + PRPSAP1-V5-BromoTag dual differential degron cell lines. **(B)** Schematic showing the selective induction of PRPS1-HA -dTAG or PRPSAP1-V5-BromoTag proteasomal degradation in response to dTAG47 or AGB1 respectively *(top)*. The effect of PRPS1 or PRPSAP1 selective degradation on PRPS1-NASP and PRPSAP1-HAT1 interaction *(bottom)*. **(C)** Pronase activity on new H3 or H4 as substrates purified from cells with or without PRPS1 depletion **(D-E)** Dox-inducible new FLAG-histone H3.3 *(C)* or H4 *(D)* interaction with histone chaperones following selective depletion of PRPS1 or PRPSAP1 for 24 hours in soluble fraction of HEK293T cells with PRPS1-HA-dTAG + PRPSAP1-V5-BromoTag dual differential degron.
